# Population modeling with machine learning can enhance measures of mental health

**DOI:** 10.1101/2020.08.25.266536

**Authors:** Kamalaker Dadi, Gaël Varoquaux, Josselin Houenou, Danilo Bzdok, Bertrand Thirion, Denis Engemann

## Abstract

**Background:** Biological aging is revealed by physical measures, *e*.*g*., DNA probes or brain scans. Instead, individual differences in mental function are explained by psychological constructs, e.g., intelligence or neuroticism. These constructs are typically assessed by tailored neuropsychological tests that build on expert judgement and require careful interpretation. Could machine learning on large samples from the general population be used to build proxy measures of these constructs that do not require human intervention?

**Results:** Here, we built proxy measures by applying machine learning on multimodal MR images and rich sociodemographic information from the largest biomedical cohort to date: the UK Biobank. Objective model comparisons revealed that all proxies captured the target constructs and were as useful, and sometimes more useful than the original measures for characterizing real-world health behavior (sleep, exercise, tobacco, alcohol consumption). We observed this complementarity of proxy measures and original measures when modeling from brain signals or sociodemographic data, capturing multiple health-related constructs.

**Conclusions:** Population modeling with machine learning can derive measures of mental health from brain signals and questionnaire data, which may complement or even substitute for psychometric assessments in clinical populations.

**Key Points:**

- We applied machine learning on more than 10.000 individuals from the general population to define empirical approximations of health-related psychological measures that do not require human judgment.
- We found that machine-learning enriched the given psychological measures via approximation from brain and sociodemographic data: Resulting proxy measures related as well or better to real-world health behavior than the original measures.
- Model comparisons showed that sociodemographic information contributed most to characterizing psychological traits beyond aging.

## Background

Quantitative measures of mental health remain challenging despite substantial efforts [1]. The field has struggled with unstable diagnostic systems [2], small sample sizes [3], and reliance on case-control studies [4]. Perhaps most importantly, mental health cannot be measured the same way diabetes can be assessed through plasma levels of insulin or glucose. Psychological constructs, *e*.*g*., intelligence or anxiety, can only be probed indirectly through lengthy expert-built questionnaires or structured examinations by a specialist. Though questionnaires often remain the best accessible option, their capacity to measure a construct is limited [5]. In practice, as full neuropsychological evaluation is not automated process but relies on expert judgement to confront multiple answers and interpret them in the context of the broader picture, such as cultural background of the participant. While the field of psychometrics has thoroughly studied the validity of psychological constructs and their measure [6, 7, 8], the advent of new biophysical measurements of the brain brings new promises [9, 10, 11]. The growth of biobanks and advances in machine learning open the door to large-scale validation of psychological measures for mental health research [12], and the hope to develop more generalizable models [13]. Yet, to be reliable, machine learning needs large labeled datasets [14]. Its application to learn imaging biomarkers of mental disorders is limited by the availability of large cohorts with high-quality neuropsychiatric diagnosis [15].

By comparison, it is easier to collect data on the general population without information on clinical conditions. For brain health, such data has lead to developing proxy measures that quantify biological aging [16, 17, 18, 11, 19, 20, 21, 22]. One counterintuitive aspect of the methodology is that measures of biological aging can be obtained by focusing on the age of a person, which is known in advance and in itself not interesting. Yet, by (imperfectly) predicting the age from brain data, machine-learning can capture the relevant signal. Based on a population of brain images, it extracts the *best guess* for the age of a person, indirectly positioning that person within the population. Individual-specific prediction errors therefore reflect deviations from what is statistically expected [23]. The brain of a person can look similar to the brains commonly seen in older (or younger) people. The resulting brain-predicted age reflects physical and cognitive impairment in adults [24, 17, 16] and reveals neurodegenerative processes [22, 25]. Can this strategy of biomarker-like proxy measures be extended to other targets beyond the construct of aging? Extrapolating from these successes, we propose to build upon large datasets to extend the collection of health-related *proxy measures*, probing mental traits. For this end, we focused on constructs fundamentally different in terms of content and methodology.

One high-stake target is intelligence, which is measured through socially administered tests and is one of the most extensively studied constructs in psychology. Fluid intelligence refers to the putatively culture-free, heritable and physiological component of intelligence [26, 27] and is a latent construct designed to capture individual differences in cognitive capacity. It has been robustly associated with neuronal maturation and is typically reflected in cognitive-processing speed and workingmemory capacity [28]. Applied to psychiatric disorders, it may help characterize psychosis, bipolar disorder, and substance abuse [29, 30].

Neuroticism is a second promising target. As a key representative of the extensively studied Big Five personality inventory, neuroticism has a long-standing tradition in the psychology of individual differences [31, 32]. Neuroticism is measured using self-assessment questionnaires and conceptualized as capturing dispositional negative emotionality including anxiety and depressiveness [33]. It has been inter-culturally validated [26, 34] and population-genetics studies have repeatedly linked neuroticism to shared genes [35, 36, 37]. Neuroticism was shown useful in psychometric screening and supports predicting real-world behavior [38, 39].

Despite strong population-level heritability [40, 41], the link between psychological constructs, brain function and genetics is still being actively researched [42, 33]. Empowered by emerging large-scale datasets, current attempts to predict fluid intelligence or neuroticism from thousands of MRI scans argue in favor of heterogeneity and weakly generalizing effects [43, 44]. This stands in contrast to the remarkable performance obtained when predicting psychometric data from language-based inputs captured by Twitter and Facebook user data [45, 46]. As MRI acquisitions can be difficult to come by in certain populations, the promises of social-media data are appealing. However, such data may lead to measurement and selection biases difficult to control. Instead, background sociodemographic data may provide an easily accessible alternative for contextualizing the heterogeneity of psychological traits [47].

Another challenge is that psychological traits are often measured using arbitrary non-physical units, *e*.*g*. education degree or monthly income. In fact, society treats individual differences as categorical or continuous, depending on the practical context. While personality has been proposed to span a continuum [48], psychiatrists treat certain people as patients and not others [49]. Therefore, a measure that performs globally poorly at a continuous scale can be sufficient to distinguish subgroups as it may be informative around the boundary region between certain classes, *e*.*g*., pilots who should fly and who should not. Choosing the granularity with which to gauge psychological constructs is difficult.

Confronting the promises of population phenotyping with the challenges of measuring psychological traits raises the following questions: 1) Can the success of brain age at characterizing health be extended to other proxy measures directly targeting mental constructs? 2) How well can various constructs related to mental health be approximated from general-purpose inputs not designed to measure specific latent constructs? 3) What is the relative merit of brain imaging and sociodemographics? We tackled these questions by using machine learning to craft *proxy measures* in order to approximate well-characterized *target measures* from brain-imaging and sociodemographic data. We studied age, fluid intelligence, and neuroticism. These targets have been, traditionally, considered as proxies for mental health and are fundamentally different in terms of scope and nature. Our results suggest that, as with brain age, proxy measures can bring value for the study of mental health that goes beyond approximating an available measure.

The paper is organized as follows: We first present a summary of the methodology and the workflow of building distinct proxy measures for age, fluid intelligence and neuroticism using machine learning (Figure 1). We then benchmark the proxy and the original target measures against real-world patterns of health-relevant behavior. Subsequently, through systematic model comparisons, we assess the relative contributions of brain imaging and sociodemographic data for prediction performance in the regression and classification settings. The complementarity between the proxy measures is, finally, discussed in the light of statistical considerations, potential data generating mechanisms, and applications for public health and clinical research.

**Figure 1.**
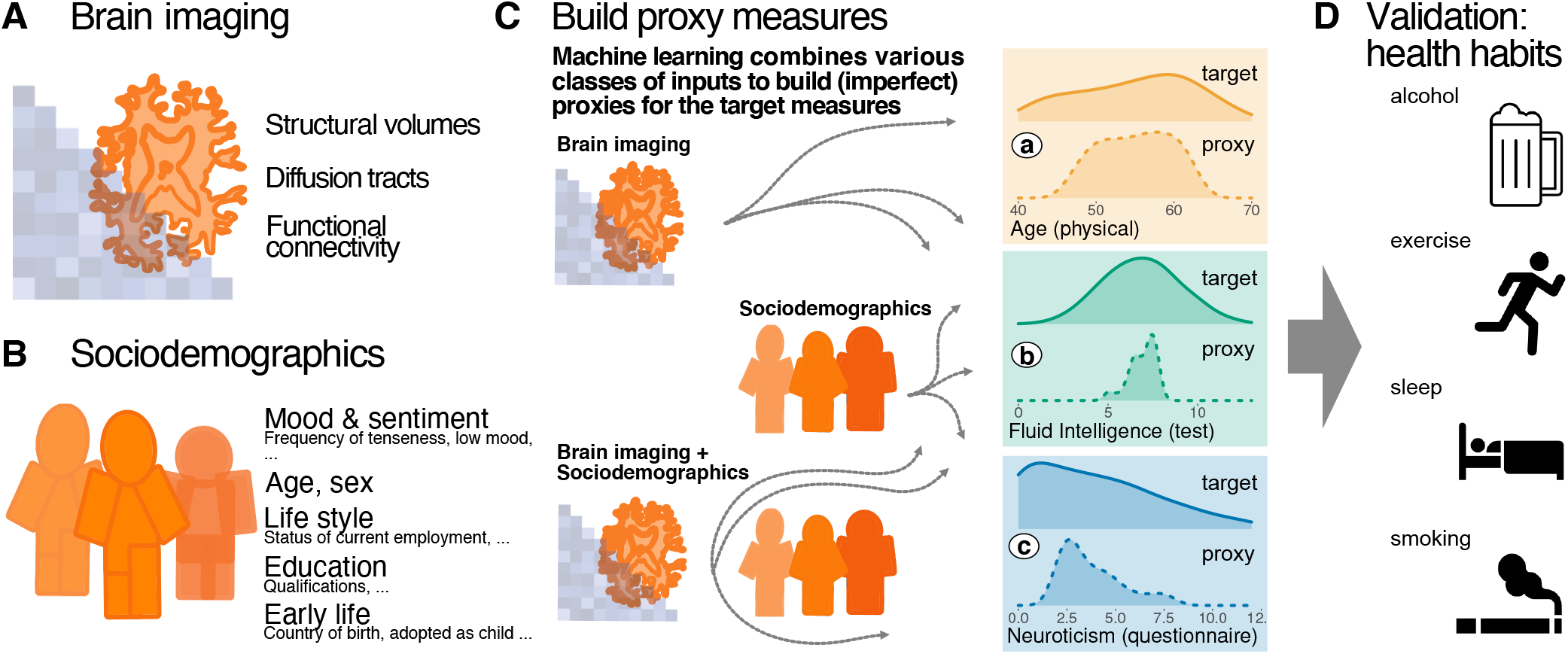
Methods workflow: building and evaluating proxy measures. We combined multiple brain-imaging modalities (**A**) with sociodemographic data (**B**) to approximate health-related biomedical and psychological constructs (**C**), *i*.*e*., brain age (accessed through prediction of chronological age), cognitive capacity (accessed through a fluid-intelligence test) and the tendency to report negative emotions (accessed through a neuroticism questionnaire). We included the imaging data from the 10 000-subjects release of the UK biobank. Among imaging data (**A**) we considered features related to cortical and subcortical volumes, functional connectivity from rfMRI based on ICA networks, and white-matter molecular tracts from diffusive directions (see Table 1 for an overview about the multiple brain-imaging modalities). We then grouped the sociodemographic data (**B**) into five different blocks of variables related to self-reported mood & sentiment, primary demographics, lifestyle, education, and early-life events (Table 2 lists the number of variables in each block). We systematically compared the approximations of all three targets based on either brain images and sociodemographics in isolation or combined (**C**) to evaluate the relative contribution of these distinct inputs. Note that proxy measures can only add to the target measures if they are not identical, *i*.*e*., if the approximation of the target from the given inputs is imperfect (guaranteed in our context as the exact data generating mechanism is unknown and causally important variables remain unobserved). Using the full model (brain imaging + sociodemographics), we benchmarked complementarity of the proxy measures and the target measures with regard to real-world patterns of health behavior (**D**), i.e., the number of alcoholic beverages, exercise (metabolic equivalent task), sleep duration and the number of cigarettes smoked. Potentially additive effects between proxies and targets were gauged using multiple linear regression. Models were developed on 50% of the data (randomly drawn) based on random forest regression guided by Monte Carlo cross-validation with 100 splits (see section Model Development and Generalization Testing). We assessed generalization and health implications using the other 50% of the data as fully independent out-of-sample evaluations (see section Statistical Analysis). Learning curves suggested that this split-half approach provided sufficient data for model construction (Figure 1 – Figure supplement 1).

## Results: validity of proxy measures

### Complementing the original measures at characterizing real-life health-related habits

To approximate age, fluid intelligence and neuroticism, we applied random-forest regression on sociodemographic data and brain images. The data was split into *validation data* for model construction (see section Model Development and Generalization Testing) and *generalization data* for statistical inference on out-of-sample predictions with independent data (see section Statistical Analysis). Our findings suggested that some information on psychological constructs can be assembled from general inputs not specifically tailored to measure these constructs, such as brain images and sociodemographic variables. The resulting proxy measures may be regarded as crude approximations of the psychological measures, but they can nonetheless capture essential aspects of the target constructs. To probe the external validity of the proxy measures, we investigated their link with real-world behavior, e.g., sleep, physical exercise, alcohol and tobacco consumption on left-out data. To probe the external validity of the proxy-measures, we investigated their link with real-world behavior, *e*.*g*., sleep, physical exercise, alcohol and tobacco consumption on left-out data. To relate such health behaviors to our proxy measures, we modeled them separately as weighted sums of predicted brain-age delta, fluid intelligence and neuroticism using multiple linear regression (section Statistical Analysis). To avoid circularity, we used the out-of-sample predictions for all proxy measures (section Model Development and Generalization Testing).

The estimated regression coefficients (partial correlations), revealed complementary associations between the proxy measures and health-related behavior (Figure 2). Similar patterns arise when considering proxy measures in isolation (Figure 2 – Figure supplement 1). Compared to other proxy measures, elevated brain-age delta was associated with increased alcohol consumption (Figure 2, first row). Levels of physical exercise were consistently associated with all three predicted targets, suggesting additive effects (Figure 2, second row). For fluid intelligence, this result, counter-intuitive from the health standpoint, could imply that higher test scores reveal a more sedentary life style. Increased sleep duration consistently went along with elevated brain age delta, but lower levels of predicted neuroticism (Figure 2, third row). This may seem counterintuitive, but is conditional on neuroticism showing a negative link with sleep duration. No consistent effect emerged for fluid intelligence. Numbers of cigarettes smoked was independently associated with all predicted targets (Figure 2, last row): Intensified smoking went along with elevated brain age delta and neuroticism but lower fluid intelligence.

**Figure 2.**
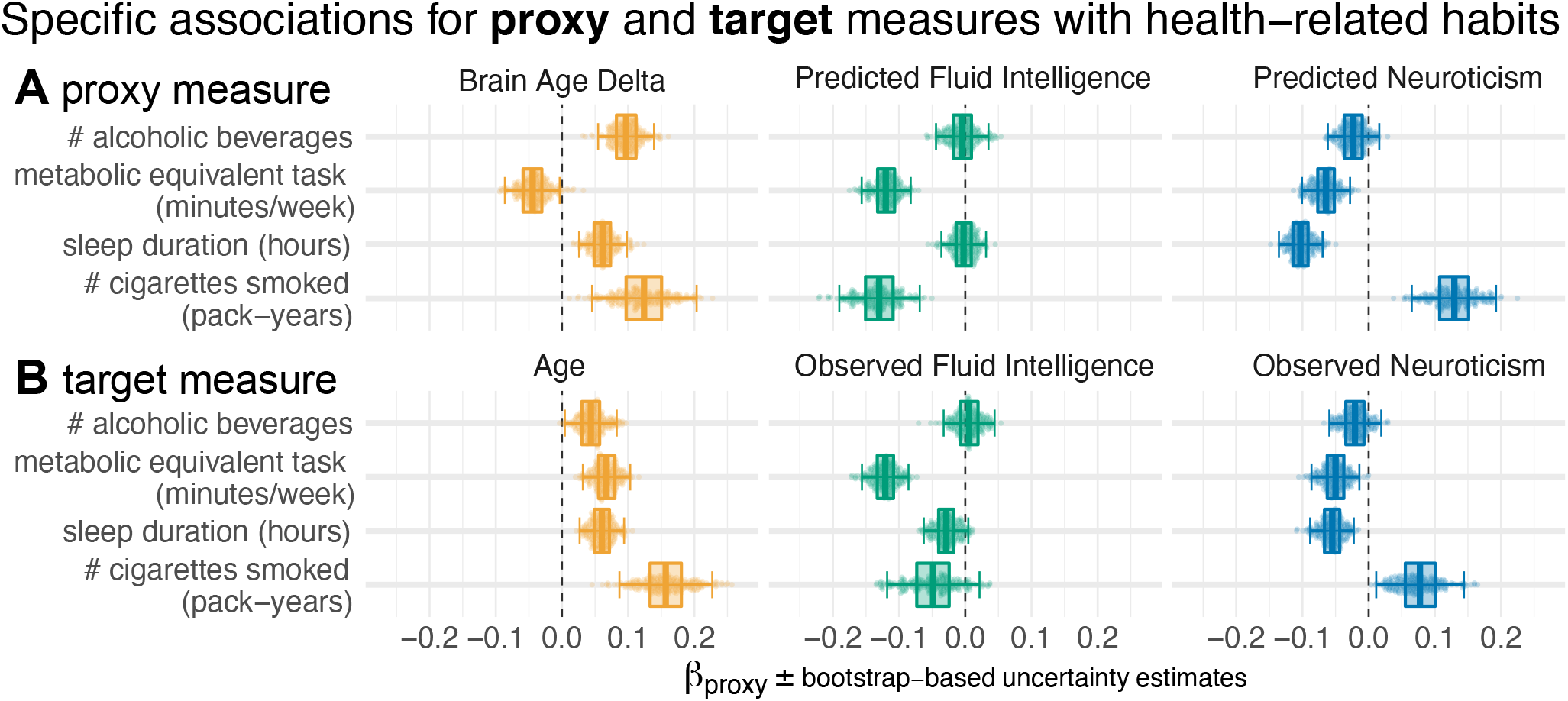
Proxy measures show systematic and complementary out-of-sample associations with health-related habits. We probed the external validity of all three proxy measures (brain age, fluid intelligence, neuroticism) based on a combination of brain images and all sociodemographic factors (see **Figure 1** for details). We investigated their out-of-sample associations with ecological indicators of mental health (sleep duration, time spent with physical exercise, number of alcoholic beverages and cigarettes consumed). To tease apart complementary and redundant effects, we constructed multiple linear regression models on out-of-sample predictions combining all three proxy measures **(A)**. For comparison, we repeated the analysis using the actual target measures **(B)** observed on the held-out data. Regression models are depicted rows-wise. Box plots summarize the uncertainty distribution of target-specific (color) regression coefficients with whiskers indicating two-sided 95% uncertainty intervals (parametric bootstrap). Dots illustrate a random subset of 200 out of 10 000 coefficient draws. The average coefficient estimate is annotated for convenience. At least two distinct patterns emerged: either the health outcome was specifically associated with one proxy measures (brain age delta and number of alcoholic beverages) or multiple measures showed additive associations with the outcome (*e*.*g*. number of pack years smoked). For target measures **(B)**, associations with health habits were often noisier or less pronounced compared to the target measures **(A)** and even a a change in direction was observed for brain age and metabolic activity. Figure 2 – Figure supplement 1 shows highly similar trends with marginal associations between proxy measures and health-related habits. Our results suggest that the proxy measures capture well health-related habits, potentially better than the original target measures, and in a complementary way across the three measures. The same patterns emerged as brain-predicted age rather than the brain age delta is used as a proxy measure (Figure 2 – Figure supplement 2). As proxy-specific deconfounding is applied, this pattern is preserved (Figure 2 – Figure supplement 3). Modeling of health-related habits jointly from proxy and target measures simultaneously revealed specific complementari. ty between proxy and target measures across multiple domains *i*.*e*. age, fluid intelligence, neuroticism (Figure 2 – Figure supplement 4).

The three proxy measures are difficult to compare on an equal footing as a delta was considered for brain age only (the difference between predicted and actual age) and aging-specific deconfounding was applied. The brain-age delta is indeed the standard practice, theoretically justified as age is on a metric scale [50] for which the difference between the predicted and the measured value has a clear meaning. Such a difference is less obvious for variables with ordinal scales as implied by psychometric measures. Second, age has a pervasive influence on virtually any biomedical entity, which motivates controlling for its effect on proxy measures. To rule out that differences in proxy measures’ associations to health-related behavior are driven by this methodological asymmetry, we repeated the main analysis from Figure 2, first, using the predicted age without computing the delta (Figure 2 – Figure supplement 2) and, second, introducing additional deconfounders for fluid intelligence and neuroticism (Figure 2 – Figure supplement 3). The resulting patterns were virtually unchanged, confirming that interpretations are robust.

A question that remains is whether the proxy measures bring additional value compared to the original target measures they were derived from. These original target measures showed similar associations to health behavior, with the same signs in most cases (Figure 2, B). At the same time, the ensuing patterns were more noisy, suggesting that empirically derived proxy measures yielded enhanced associations with health behavior. This inference may be difficult as differences between targets and proxies were not always easy to pinpoint visually. To implement a more rigorous statistical approach, we built comprehensive models of each respective health-related habit in which we used all proxies (predicted age, predicted fluid intelligence, predicted neuroticism) and all targets (age, fluid intelligence, neuroticism) simultaneously as predictors (Figure 2 – Figure supplement 4). The results show systematic additive effects of proxies and targets across the three target domains and the four health-habits. These trends are well-captured by the hypothesis tests of the respective linear models (Table S3). As targets and proxies may be systematically intercorrelated, multicollinearity may corrupt these inferences. Inspection of variance inflation factors (VIF)— a measure that reveals how well a given predictor can be approximated by a linear combination of the other predictors— argued in favor of low to moderate levels of multicollinearity (Table S4). Indeed, all VIF values fell between 3 and 1, whereas, classically, values above 5 or 10 are considered as thresholds [51] for pathological collinearity. This suggests that the model inferences are statistically sound.

### The relative importance of brain and sociodemographic data depends on the target

In a second step, we investigated the relative performance of proxy measures built from brain signals and distinct sociodemographic factors for the three targets: age, fluid intelligence and neuroticism. Among the sociodemographic variables there was one block for each target explaining most of the prediction performance (Figure 3, dotted outlines). Combining all sociodemographic variables did not lead to obvious enhancements (Figure 3 – Figure supplement 2). For age prediction, variables related to current life-style showed by far the highest performance. For fluid intelligence, education performed by far best. For neuroticism, mood & sentiment clearly showed the strongest performance.

**Figure 3.**
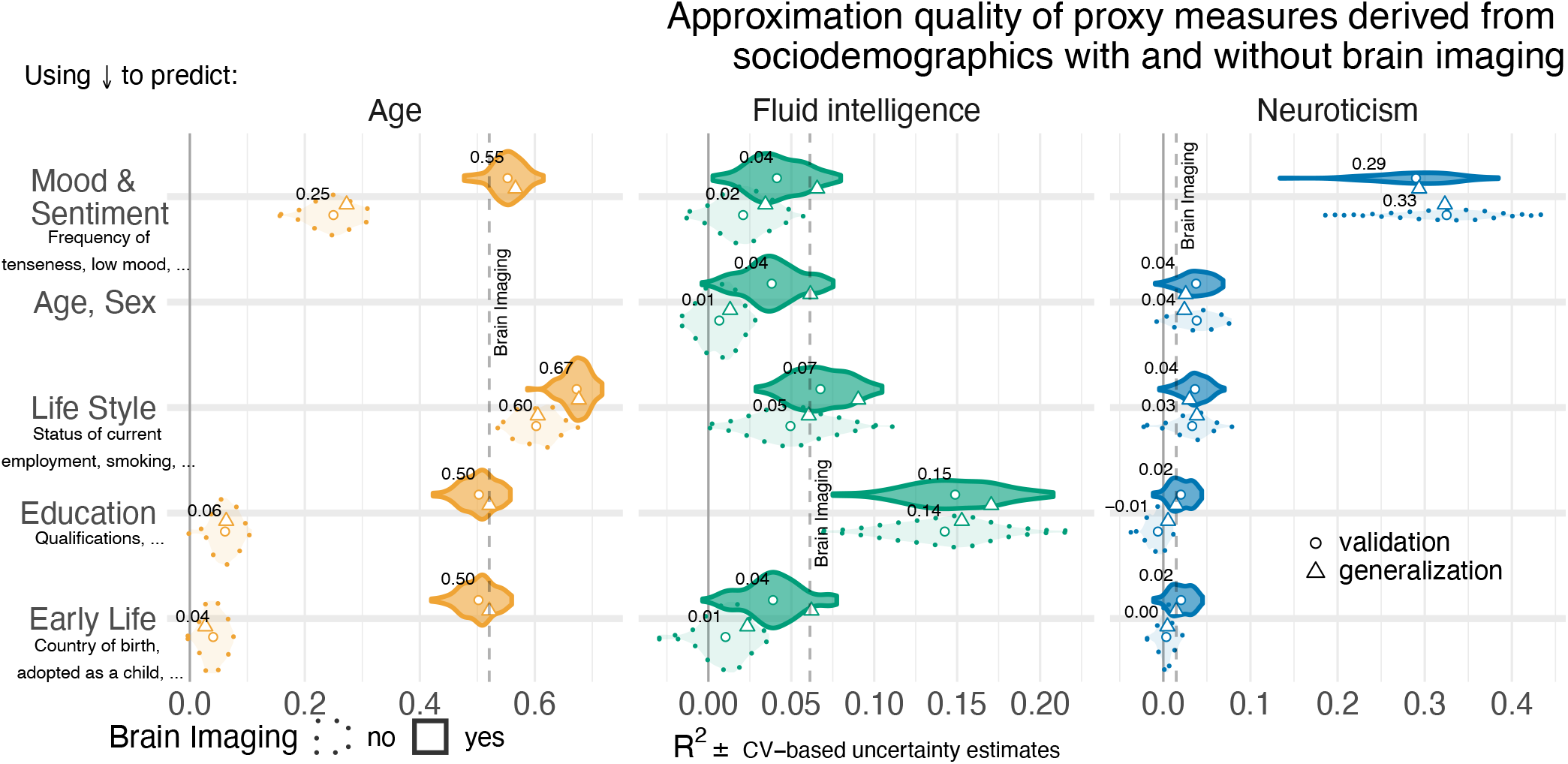
Approximation performance of proxy measures derived from sociodemographic data and MRI. We report the *R*^2^ metric to facilitate comparisons across prediction targets. The cross-validation (CV) distribution (100 Monte Carlo splits) on the validation dataset is depicted by violins. Drawing style indicates whether brain imaging (solid outlines of violins) was included in addition or not (dotted outlines of violins). Dots depict the average performance on the validation data across CV-splits. Pyramids depict the performance of the average prediction (CV-bagging) on held-out generalization datasets. For convenience, the mean performance on the validation set is annotated for each plot. Vertical dotted lines indicate the average performance of the full MRI model. The validation and held-out datasets gave similar picture of approximation performance with no evidence for cross-validation bias [52]. For the averaged out-of-sample predictions, the probability of the observed performance under the null-distribution and the uncertainty of effect sizes were formally probed using permutation tests and bootstrap-based confidence intervals (Table S1). Corresponding statistics for the baseline performance of models solely based on brain imaging (vertical dotted lines) are presented in Table S5. Figure 3 – Figure supplement 1 shows approximation results based on MRI. Figure 3 – Figure supplement 2 presents results based on all sociodemographic factors.

Combining MRI and sociodemographics, enhanced age prediction systematically on all four blocks of variables (Figure 3 solid outlines, and Table S1). The benefit of brain-imaging features was less marked for prediction of fluid intelligence or neuroticism. With fluid intelligence, brain-imaging data improved the performance statistically significantly for all models, yet, with small effect sizes (Table S1). For neuroticism, no systematic benefit of including brain images alongside sociodemographics emerged (Table S1, bottom row). Nevertheless, brain data was sufficient for statistically significant approximation of the target measures in all three targets (Table S5).

Psychological measures often come without physical scales and units [50]. In practice, clinicians and educators use them with specific thresholds for decision making. To investigate empirically-defined proxy measures beyond continuous regression, we performed binary classification of extreme groups obtained from discretizing the targets using the 33_rd_ and 66_th_ percentiles, following the recommendations by Gelman and Hill 2006 regarding discrete variable encoding strategies. Furthermore, we measured accuracy with the area under the classification accuracy curve (AUC) which is only sensitive to ranking, ignoring the scale of the error. Classification performance visibly exceeded the chance level (AUC > 0.5) for all models (Figure 4) and approached or exceeded levels considered practically useful (AUC > 0.8) [49]. Across proxy measures, models including sociodemographics performed best but the difference between purely sociodemographic and brain-based models was comparably weak, at the order of 0.01-0.02 AUC points (Table S2). Using brain data only led to worse performance, yet, still better than chance as revealed by permutation testing (Table S6).

**Figure 4.**
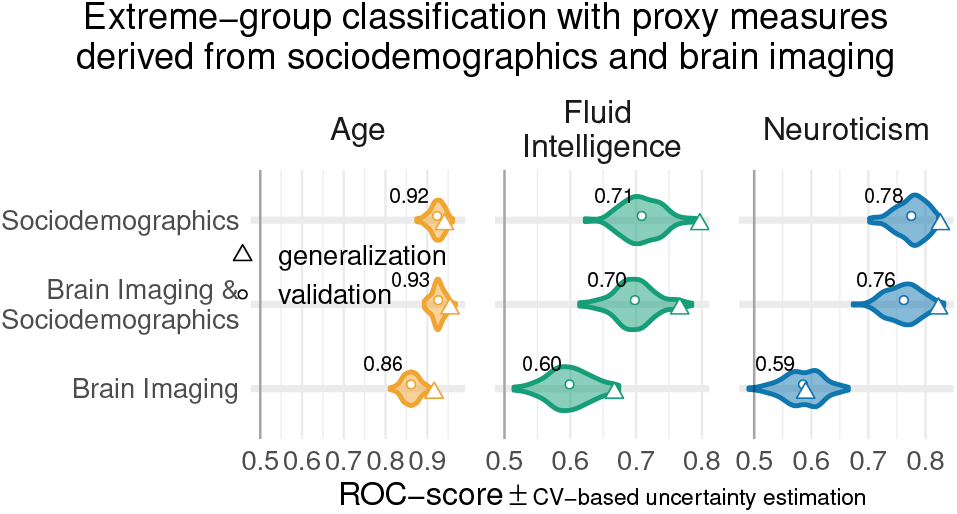
Classification analysis from imaging, sociodemographics and combination of both data. For classification of extreme groups instead of continuous regression, we split the data into low vs high groups based on 33_rd_ and 66_th_ percentiles. Visual conventions follow Figure 3. We report the accuracy in AUC. Models including sociodemographics performed visibly better than models purely based on brain imaging. Differences between brain-imaging and sociodemographics appeared less pronounced as compared to the fully-fledged regression analysis. For the average out-of-sample predictions, the probability of the observed performance under the null-distribution and the uncertainty of effect sizes were formally probed using permutation tests and bootstrap-based confidence intervals (Table S2). Corresponding statistics for the baseline performance of models solely based on brain imaging (vertical dotted lines) are presented in Table S6. Overall, when moving from the more difficult full-scale regression problem to extreme-group classification problem with purely ranking-based scores, the relative differences between brain-based and sociodemographics-based prediction gradually faded away.

## Discussion

Guided by machine learning, we empirically derived proxy measures that combine multiple sources of information to capture extensively validated target measures from psychology. These proxy measures all showed complementary associations with real-world health indicators beyond the original targets. The combination of brain imaging and target-specific sociodemographic inputs often improved approximation performance.

### Empirically-derived proxy measures: validity and practical utility

In our study, construct validity [6, 54, 7] of the corresponding proxy measures was supported by the gain in prediction performance brought by specific sociodemographic factors (Figure 3). Association with health-relevant habits added external validity to the proxy measures (Figure 2). The complementary patterns related to traditional construct semantics: High consumption of cigarettes is associated with neuroticism [55], excessive drinking may lead to brain atrophy and cognitive decline [56] – both common correlates of elevated brain age [22, 57].

Can our empirically-derived proxy measures, thus, substitute for specific psychometric instruments? A mental-health professional may still prefer an established routine for clinical assessment, relying on interviews and personality questionnaires with implicit experience-based thresholds. Inclusion of brain imaging may even seem to yield diminishing returns when approximating high-level psychological traits. Yet, it could simply be a matter of time until more effective acquisition protocols will be discovered alongside useful signal representations. Including brain imaging, rather seems a “safe bet” as machine learning is often capable of selecting relevant inputs [11, 58] and costs of MRI-acquisition can be amortized by clinical usage. Empirically-derived proxy measures may open new doors where tailored assessment of latent constructs is not applicable due to lack of specialized mental-health workforce or sheer cost.

### Constructs of mental-health can be accessed from general-purpose data

Brain age has served as landmark in this study. It has been arguably the most discussed candidate for a surrogate biomarker in the brain imaging literature [16, 17, 24]. With mean absolute errors around 4 years, up to 67% variance explained, and AUC-scores up to 0.93 in the classification setting, our results compare favorably to the recent brain-age literature within the UK Biobank [19, 59] and in other datasets [22, 11], though we relied on off-the-shelf methods and not custom deep learning methods [60]. Applying the same approach to psychological constructs (fluid intelligence, neuroticism), we found that approximation from brain imaging data or sociodemographic descriptors was generally harder.

It is important to recapitulate that approximation quality on these differently measured targets has a different meaning. Age is measured with meaningful physical units (years) on a ratio scale [50] (Selma is *twice as old* as Bob). Psychometric scores are unit-free, which may provoke ambiguity regarding the level of measurement [54]. Their implied scales may be considered as interval (the *difference between* Bob’s and Selma’s intelligence is -0.1 standard deviations) if not ordinal (Bob’s intelligence was *ranked below* Selma’s) [50]. In day-to-day psychological practice, these scores are often used via practically-defined thresholds, *e*.*g*. school admission or pilot candidate selection [61, 62]. In the classification setting, all proxy measures approached or exceeded a performance of 0.80 deemed relevant in biomarker development [49], though to be fair, they approximated established psychometric targets (proxy measures themselves) and not a medical condition. Different proxy measures should, thus, be subjected to different standards, depending on the granularity of the implied measurement scale.

A more complete view on how the proxy measures capture mental-health constructs emerges from their associations with real-world behavior (Figure 2). Indeed, the associations with proxy measures (Figure 2 **B**) were less noisy and more consistent then with the target measures (Figure 2 **A**), regardless of their approximation quality. This may seem surprising at first, but the target measures are themselves noisy and of imperfect validity. These measures correspond to traditional tests which, in practice, must be interpreted by an expert, actively confronting their output with broader information on the individual. For instance, IQ scores are typically normalized across age groups. However, extending such a normalization approach to many factors (socio-economic status, culture, gender) poses fundamental high-dimensional statistics challenges. Conversely, using machine learning to assemble proxy measures by mapping the targets to rich sociodemographic and brain data implicitly contextualizes them. In this respect, the resulting measures capture more general signal than the original tests. Here, machine learning could be seen as mimicking the work of a mental health expert who carefully compares psychometric results with other facts known about an individual and its reference population.

### The benefits offered by brain data depend on the target construct

All brain-derived approximations were statistically meaningful. Yet, only for age prediction, imaging data by itself led to convincing performance. For fluid intelligence and neuroticism, sociodemographic factors were the most important determinants of prediction success. The best-performing sociodemographic models were based on inputs semantically close to these targets, *i*.*e*., education details or mood & sentiment. While those results support construct validity, they may come with a certain risk of circularity. The causal role of those predictors is not necessarily clear as better educational attainment is heritable itself [63] and may reinforce existing cognitive abilities. Similarly, prolonged emotional stress due to life events may exacerbate existing dispositions to experience negative emotions captured by neuroticism [64], traits which commonly help accumulate stressful life events [38]. Nevertheless, for fluid intelligence but not neuroticism, brain imaging added incremental value when combined with various sociodemographic predictors. This may suggest that the cues for neuroticism conveyed by brain imaging were already present in sociodemographic predictors, hinting at common causes. Off note, in the specific context of aging, the empirical distinction between brain age and cognitive age is reflecting a similar intuition [65].

### Limitations

Additional constructs and psychometric tools could have been visited. The broader construct of intelligence is often estimated using a general factor model with multiple correlated tests. While this is obviously useful for normative assessments, measures of fluid intelligence can also serve a situational fitness signal [30]. There is a wealth of questionnaires for measuring negative emotionality and neuroticism, specifically. Yet, we could only study the EPQ scale provided by the UK Biobank. A complementary approach would be to estimate latent factors by pooling all non-imaging data semantically related to neuroticism [66]. Here, we considered established target measures “as is”, instead of derivatives.

It terms of mental-health research, this study falls short of directly testing the clinical relevance of estimated proxy measures. Even in a very large general-population cohort such as the UK Biobank, there are only a few hundred diagnosed cases of mental disorders (ICD-10 mental-health diagnoses from the F chapter) with brain-imaging data available. As a result, we could not directly assess the performance of proxy measures in clinical populations. The low number of diagnosed mental disorders in UK Biobank highlights the practical importance of studying mental health as a continuous, in addition to diagnosed conditions. Indeed, a public health perspective calls for targeting individual differences in health, not only pathology. Psychological constructs such as IQ and neuroticsm are important factors of the epidemiology of psychiatric disorders [38, 30, 29, 67], and accelerated brain aging is associated with various neurological conditions [18, 17, 25]. Yet, few cohorts come with extensive neuropsychological testing. Validated proxies of these constructs open the door to including them in epidemiological studies as secondary outcomes or additional explanatory variables.

### Conclusion: Proxy measures may enhance the validity of constructs gauging mental health

In population studies of mental health, individual traits are captured via lengthy assessments, tailored to specific brain and psychological constructs. We have shown that proxy measures built empirically from general-purpose data can capture these constructs and can improve upon traditional measures when studying real-world health patterns. Proxy measures can make psychological constructs available to broader, more ecological studies building on large epidemiological cohorts or real-world evidence. This can make the difference where psychological constructs are central to developing treatment and prevention strategies, but direct measures have not been collected.

## Methods

To facilitate reproduction, understanding, and reuse, we have made all data analysis and visualization source code available on Github [68].

### Dataset

The United Kingdom Biobank (UKBB) database is to date the most extensive large-scale cohort aimed at studying the determinants of the health outcomes in the general adult population. The UKBB is openly accessible and has extensive data acquired on 500 000 individuals aged 40-70 years covering rich phenotypes, health-related information, brain-imaging and genetic data [12]. Participants were invited for repeated assessments, some of which included MR imaging. For instance, cognitive tests that were administered during an initial assessment were also assessed during the follow-up visits. This has enabled finding for many subjects at least one visit containing all heterogeneous input data needed to develop the proposed proxy measures. The study was conducted using the UKBB Resource Application 23827.

### Participants

All participants gave informed consent. The UKBB study was examined and approved by the North West Multi-centre Research Ethics Committee. We considered participants who have responded to cognitive tests, questionnaires, and have access to their primary demographics and brain images [69]. Out of the total size of UKBB populations, we found 11 175 participants who had repeated assessments overlapping with the first brain imaging release [70]. Note that the features (sociodemographic variables) that we included in the analysis are measures that are self-reported during a follow-up imaging visit. The demographics are 51.6% female (5 572) and 48.3% male (5 403), and an age range between 40-70 years (with a mean of 55 years and standard deviation of 7.5 years). The data for model training were selected using a randomized split-half procedure yielding 5 587 individuals. The remaining subjects were set aside as a held-out set for generalization testing (see section Model development and generalization testing). We made sure that the subjects used for model training and generalization testing were strictly non-overlapping.

Learning curves documented that the training split was sufficiently large for constructing stable prediction models Figure 1 – Figure supplement 1 with profiles of performance similar to latest benchmarks on model complexity in the UK Biobank [71]. Moreover, simulations and empirical findings suggest that larger testing sets are more effective at mitigating optimistic performance estimates [72, 52]. Together, this provided a pragmatic solution to the inference-prediction dilemma [58, 73] given the two objectives of the present investigation to obtain reasonably good predictive models, while at the same time performing parameter inference of statistical models developed on the left-out data.

To establish specific comparisons between models based on sociodemographics, brain data or their combinations we exclusively considered the cases for which MRI scans were available. The final sample sizes used for model construction and generalization testing then depended on the availability of MRI: For age and fluid intelligence, our randomized split-half procedure (see section Model development and generalization testing) yielded 4203 cases for model building and 4157 for generalization. For cases with valid neuroticism assessment, fewer brain images were available, which yielded 3550 cases for model building and 3509 for generalization.

### Data acquisition

Sociodemographic data (non-imaging) was collected with self-report measures administered through touchscreen questionnaires, complemented by verbal interviews, physical measures, biological sampling and imaging data. MRI data were acquired with the Siemens Skyra 3T using a standard Siemens 32-channel RF receiver head coil [74]. We considered three MR imaging modalities as each of them potentially captures unique neurobiological details: structural MRI (sMRI/T1), resting-state functional MRI (rs-fMRI) and diffusion MRI (dMRI). For technical details about the MR acquisition parameters, please refer to [70]. We used image-derived phenotypes (IDPs) of those distinct brain-imaging modalities, as they provide actionable summaries of the brain measurements and encourage comparability across studies.

#### Target measures

As our target measures for brain age modeling, we use an individual’s age at baseline recruitment (UKBB code “21022-0.0”). Fluid intelligence, was assessed using a cognitive battery designed to measure an individual’s capacity to solve novel problems that require logic and abstract reasoning. In the UK Biobank, the fluid intelligence test (UKBB code “20016-2.0”) comprises thirteen logic and reasoning questions that were administered via the touchscreen to record a response within two minutes for each question. Therefore, each correct answer is scored as one point with 13 points in total^1^. Neuroticism (UKBB code “20127-0.0”) was measured using a shorter version of the revised Eysenck Personality Questionnaire (EPQ-N) comprised of 12-items [32]. Neuroticism was assessed during Biobank’s baseline visit. The summary of the individual’s scores ranges from 0 to 12 that assess dispositional tendency to experience negative emotions ^2^.

In the course of this work, a question that emerged concerned the size of the gap between age at baseline recruitment and MRI-scan time and its potential impact on the analysis. Supplementary checks indicated that the age gap was at least 5 years for most participants. Yet, from a statistical perspective, the two age measures turned out highly interchangeable (Figure S2) and global conclusions remained unchanged (Figure S3).

### Sociodemographic data

In this work, we refer to non-imaging variables broadly as sociodemographics excluding the candidate targets fluid intelligence and neuroticism. To approximate latent constructs from sociodemographics, we included 86 non-imaging inputs (Table S7) which are the collection of variables reflecting each participant’s demographic and social factors *i*.*e*., sex, age, date and month of birth, body mass index, ethnicity, exposures at early life –*e*.*g*. breast feeding, maternal smoking around birth, adopted as a child– education, lifestyle-related variables –*e*.*g*. occupation, household family income, household people living at the same place, smoking habits–, and mental-health variables. All these data were self-reported. We then assigned these 86 variables to five groups based on their relationships. Based on our conceptual understanding of the variables, we name assigned them to one out of five groups: **1)** mood & sentiment, **2)** primary demographics as age, sex, **3)** lifestyle, **4)** education, **5)** early life. We then investigated the intercorrelation between all 86 variables to ensure that the proposed grouping is compatible with their empirical correlation structure Figure S1.

The sociodemographic groups had varying amounts of missing data. For *e*.*g*. the source of missingness is concerned with the participants lifestyle habits such as smoking and mental health issues [77]. To deal with this missingness in the data using imputation [78], we used column-wise replacement of missing information with the median value calculated from the known part of the variable. We subsequently included an indicator for the presence of imputed for down-stream analysis. Such imputation is well suited to predictive models [79].

### Image processing to derive phenotypes for machine learning

MRI data preprocessing were carried out by UKBB imaging team. The full technical details are described elsewhere [70, 74]. Below, we describe briefly the custom processing steps that we used on top of the already preprocessed inputs.

#### Structural MRI

This type of data analysis on T1-weighted brain images are concerned with morphometry of the gray matter areas *i*.*e*. the quantification of size, volume of brain structures and tissue types and their variations under neuropathologies or behavior [80]. For example, volume changes in gray matter areas over lifetime are associated with: brain aging [81], general intelligence [82] and brain disease [83]. Such volumes are calculated within pre-defined ROIs composed of cortical and sub-cortical structures [84] and cerebellar regions [85]. We included 157 sMRI features consisting of volume of total brain and grey matter along with brain subcortical structures^3^. All these features are pre-extracted by UKBB brain imaging team [70] and are part of data download. We concatenated all inputs alongside custom-built fMRI features for predictive analysis (feature union).

#### Diffusion weighted MRI

Diffusion MRI enables to identify white matter tracts along principal diffusive direction of water molecules, as well as the connections between different gray matter areas [88, 89]. The study of these local anatomical connections through white matter are relevant to the understanding of neuropathologies and functional organization [90]. We included 432 dMRI skeleton features of FA (fractional anisotropy), MO (tensor mode) and MD (mean diffusivity), ICVF (intra-cellular volume fraction), ISOVF (isotropic volume fraction) and OD (orientation dispersion index) modeled on many brain white matter structures extracted from neuroanatomy^4^. For extensive technical details, please refer to [92]. The skeleton features we included were from category134 shipped by the UKBB brain-imaging team and we used them without modification.

#### Functional MRI

Resting-state functional MR images capture low-frequency fluctuations in blood oxygenation that can reveal ongoing neuronal interactions in time forming distinct brain networks [93]. Functional connectivity within these brain network can be linked to clinical status [94], to behavior [70], or to psychological traits [44]. We also included resting-state connectivity features based on the time-series extracted from Independent Component Analysis (ICA) with 55 components representing various brain networks extracted on UKBB rfMRI data [70]. These included the default mode network, extended default mode network and cingulo-opercular network, executive control and attention network, visual network, and sensorimotor network. We measured functional connectivity in terms of the between-network covariance. We estimated the covariance matrices using Ledoit-Wolf shrinkage [95]. To account for the fact that covariance matrices live on a particular manifold, *i*.*e*., a curved non-Euclidean space, we used the tangent-space embedding to transform the matrices into a Euclidean space [96, 97] following recent recommendations [98, 99]. For predictive modeling, we then vectorized the covariance matrices to 1 485 features by taking the lower triangular part. These steps were performed with NiLearn [100].

### Comparing predictive models to approximate target measures

#### Imaging-based models

First, we focused on purely imaging-based models based on exhaustive combinations of the three types of MRI modalities (see Table 1 for an overview). This allowed us to study potential overlap and complementarity between the MRI-modalities. Preliminary analyses revealed that combining all MRI data gave reasonable results with no evident disadvantage over particular combinations of MRI modalities (Figure 3 – Figure supplement 1), hence, for simplicity, we only focused on the full MRI model in subsequent analyses.

**Table 1.** Imaging-based models.

| In-dex | Name | # variables | # groups |
| --- | --- | --- | --- |
| 1 | brain volumes (sMRI) | 157 | 1 |
| 2 | white matter (dMRI) | 432 | 1 |
| 3 | functional connectivity (fMRI) | 1485 | 1 |
| 4 | sMRI, dMRI | 589 | 2 |
| 5 | sMRI, fMRI | 1642 | 2 |
| 6 | dMRI, fMRI | 1917 | 2 |
| 7 | sMRI, dMRI, fMRI (full MRI) | 2074 | 3 |

#### Sociodemographic models

We composed predictive models based on non-exhaustive combinations of different types of sociodemographic variables. To investigate the relative importance of each class of sociodemographic inputs, we performed systematic model comparisons. We were particularly interested in studying the relative contributions of early-life factors as compared to factors related to more recent life events such as education as well as factors related to current circumstances such as mood & sentiment and life-style. The resulting models based on distinct groups of predictors are listed in Table 2 (for additional details see Table S7 and Figure S1).

**Table 2.** Non-imaging baseline models or sociodemographic models based on single group. Variables in each group are described at corresponding section sociodemographic data

| Index | Name | # variables |
| --- | --- | --- |
| 1 | Mood & Sentiment (MS) | 25 |
| 2 | Age, Sex (AS) | 5 |
| 3 | Life style (LS) | 45 |
| 4 | Education (EDU) | 2 |
| 5 | Early Life (EL) | 9 |

#### Combined imaging and sociodemographic models

In the next step, we were interested in how brain-related information would interact within each of these sociodemographic models. For example, information such as the age of an individual, or the level of education, may add important contextual information to brain images. We therefore considered an alternative variant for each of the models in Table 2 that included all MRI-related features (2 074 additional features) as described at section image processing to derive phenotypes for machine learning.

#### Predictive model

Linear models are recommended as default choice in neuroimaging research [98, 101] especially when datasets include fewer than 1000 data points. In this study approximated targets generated by distinct underlying mechanisms based on multiple classes of heterogenous input data with several thousands of data points. We hence chose the non-parametric random forest algorithm that can be readily applied on data of different units for non-linear regression and classification [102] with mean squared error as impurity criterion. To improve computation time we fixed tree-depth to 250 trees, a hyper-parameter that is not usually not tuned but set to a generous number as performance plateaus beyond a certain number of trees [103, ch. 15]. Preliminary analyses suggested that additional trees would not have led to substantial improvements in performance. We used nested cross-validation (5-fold grid search) to tune the depth of the trees as well as the number of variables considered for splitting (see Table 3 for a full list of hyper-parameters considered).

**Table 3.**
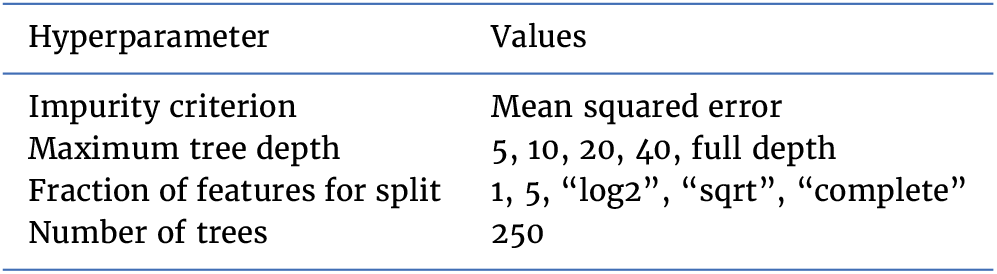
Random forest hyperparameters and tuning with grid search (5 fold cross-validation).

| Hyperparameter | Values |
| --- | --- |
| Impurity criterion | Mean squared error |
| Maximum tree depth | 5, 10, 20, 40, full depth |
| Fraction of features for split | 1, 5, "log2", "sqrt", "complete" |
| Number of trees | 250 |

#### Classification analysis

We also performed classification analysis on the continuous targets. Adapting recommendations from Gelman and Hill [53], we performed discrete variable encoding of the targets leading to extreme groups based on the 33_rd_ and 66_th_ percentiles (see Table 4 for the number of classification samples per group). This choice avoids including samples near the average outcome for which the input data may be indistinct. We were particularly interested in understanding whether model performance would increase when moving toward classifying extreme groups. For this analysis, we considered all three types of models (full MRI 2074 features from imaging-based models, all sociodemographics variables, total 86 variables see section, combination of full MRI and all sociodemographics, a total 2160 variables see section (See section Comparing predictive models to approximate target measures). When predicting age, we excluded the age & sex sociodemographic block from all sociodemographic variables which then yielded a total of 81 variables. To assess the performance for classification analysis, we used the area under the curve (AUC) of the receiver operator characteristic (ROC) as an evaluation metric [101].

**Table 4.**
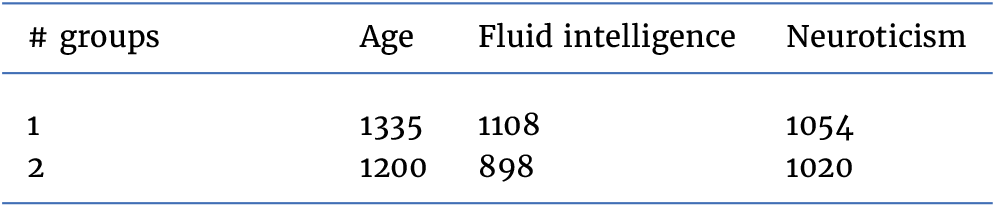
Number of samples for classification analysis (N).

| # groups | Age | Fluid intelligence | Neuroticism |
| --- | --- | --- | --- |
| 1 | 1335 | 1108 | 1054 |
| 2 | 1200 | 898 | 1020 |

### Model development and generalization testing

Before any empirical work, we generated two random partitions of the data, one validation dataset for model construction and one held-out generalization dataset for studying out-of-sample associations using classical statistical analyses.

For cross-validation, we then subdivided the validation set into 100 training- and testing splits following the Monte Carlo resampling scheme (also referred to as shuffle-split) with 10% of the data used for testing. To compare model performances based on paired tests, we used the same splits across all models. Split-wise testing performance was extracted and carried forward for informal inference using violin plots (Figure 3,Figure 4). For generalization testing, predictions on the held-out data were generated from all 100 models from each cross-validation split.

On the held-out set, unique subject-wise predictions were obtained by averaging across folds and occasional duplicate predictions due to Monte Carlo sampling which could produce multiple predictions per subject^5^. Such strategy is known as CV-bagging [104, 105] and can improve both performance and stability of results^6^. The resulting averages were reported as point estimates in Figures 3,4, and 3 – Figure supplement 1 and used as proxy measures in the analysis of health-related behaviors Figure 2.

### Statistical analysis

#### Resampling statistics for model comparisons on the held-out data

To assess the statistical significance of the observed model performance and the differences in performance between the models, we computed resampling statistics of the performance metrics on the held-out generalization data not used for model construction [106]. Once unique subject-wise predictions were obtained on the held-out generalization data by averaging the predictions emanating from each fold of the validation set (cv-bagging), we computed null- and bootstrap-distributions of the observed test statistic on the held-out data, i.e., *R*^2^ score for regression and *AUC* score for classification.

#### Baseline comparisons

To obtain a p-value for baseline comparisons (*could the prediction performance of a given model be explained by chance?*) on the held-out data, we permuted targets 10 000 times and then recomputed the test statistic in each iteration. P-values were then defined as the probability of the test statistic under null distribution being larger than the observed test statistic. To compute uncertainty intervals, we used bootstrap, recomputing the test statistic after resampling 10 000 times with replacement and reporting the 2.5 and 97.5 percentiles of the resulting distribution.

#### Pairwise comparisons between models

For model comparisons, we considered the out-of-sample difference in *R*^2^ or *AUC* between any two models. To obtain a p-value for model comparisons (*could the difference in prediction performance between two given models be explained chance?*) on the held-out data, we permuted the scores predicted by model A and model B for every single prediction 10 000 times and then recomputed the test statistic in each iteration. We omitted all cases for which only predictions from one of the models under comparison was present. P-values were then defined as the probability of the absolute of the test statistic under null distribution being larger than the absolute observed test statistic. The absolute was considered to account for differences in both directions. Uncertainty intervals were obtained from computing the 2.5 and 97.5 percentiles of the bootstrap distribution based on 10 000 iterations. Here, predictions from model A and model B were resampled using identical resampling indices to ensure a meaningful paired difference.

#### Out-of-sample association between proxy measures and health-related habits

##### Computation of brain age delta and de-confounding

For association with health-contributing habits (Table 5), we computed the brain age delta as the difference between predicted age and actual age:

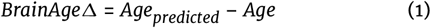

**Table 5.**
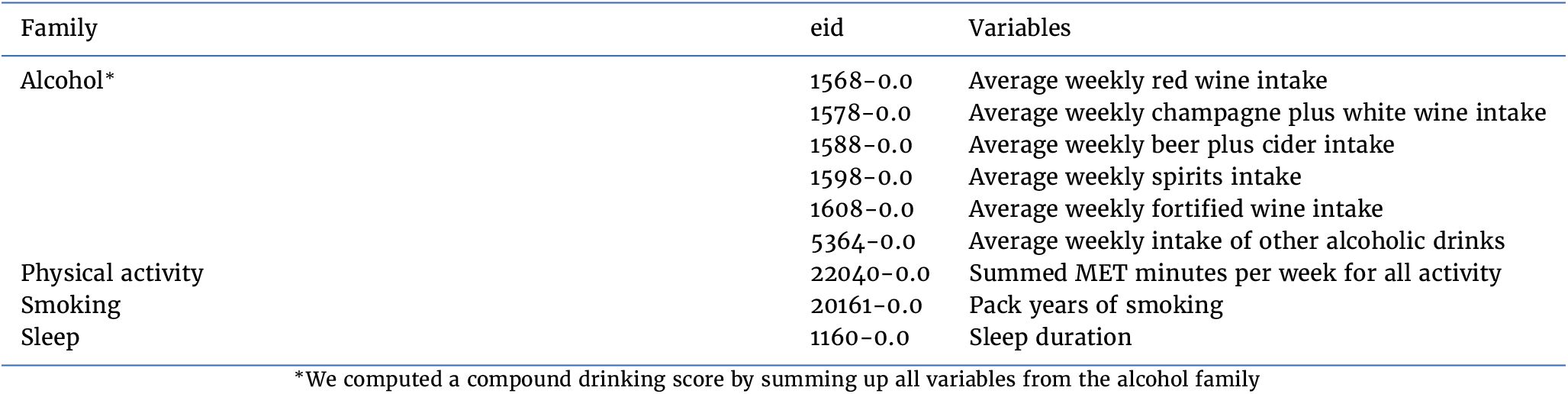
Extra health variables used for correlation analysis with subject-specific predicted scores.

As age prediction is rarely perfect, the residuals will still contain age-related variance which commonly leads to brain age bias when relating the brain age to an outcome of interest, *e*.*g*., sleep duration [107]. To mitigate leakage of age-related information into the statistical models, we employed a de-confounding procedure in line with [108] and [11, eqs. 6-8] consisting in residualizing a measure of interest (*e*.*g*. sleep duration) with regard to age through multiple regression with quadratic terms for age. To minimize computation on the held-out data, we first trained a model relating the score of interest to age on the validation set to then derive a de-confounding predictor for the held-out generalization data. The resulting de-confounding procedure for variables in the held-out data amounts to computing an age-residualized predictor *measure*_*resid*_ from the measure of interest (*e*.*g*. sleep duration) by applying the following quadratic fit on the validation data:

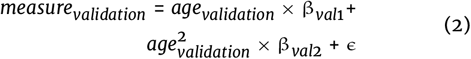

The de-confounding predictor was then obtained by evaluating the weights β_*val*1_ and β_*val*2_ obtained from Equation 2 on the generalization data:

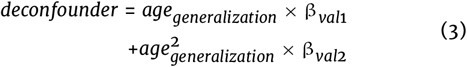

We performed this procedure for all target measures, to study associations not driven by the effect of age. For supplementary analyses presented in figure Figure 2 – Figure supplement 3, the same procedure was applied, substituting age for fluid intelligence and neuroticism, respectively.

##### Health-related habits regression

We then investigated the joint association between proxy measures of interest and health-related habits (Table 5) using multiple linear regression. For simplicity, we combined all brain imaging and all sociodemographics variables (Figure 3, Figure 3 – Figure supplement 1, Figure 3 – Figure supplement 2). The ensuing model can be denoted as

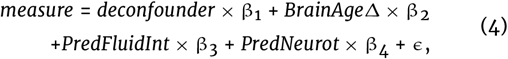

where *deconfounder* is given by Equation 2. Prior to model fitting, rows with missing inputs were omitted. For comparability, we then applied standard scaling on all outcomes and all predictors.

The parametric bootstrap was a natural choice for uncertainty estimation, as we used standard multiple linear regression which provides a well defined procedure for mathematically quantifying its implied probabilistic model. Computation was carried out using sim function from the arm package as described in [53, Ch.7,pp.142-143]. This procedure can be intuitively regarded as yielding draws from the posterior distribution of the multiple linear regression model under the assumption of a uniform prior. For consistency with previous analyses, we computed 10000 draws.

For supplementary analysis in Figure 2 – Figure supplement 2, the brain-predicted age instead of the delta was used:

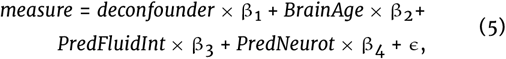

For supplementary analysis in Figure 2 – Figure supplement 3, additional deconfounders were introduced.

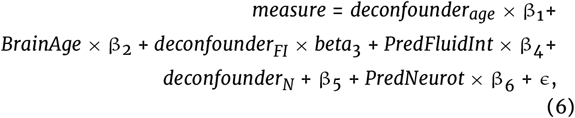

where *deconfounder*_*FI*_ is the deconfounder for fluid intelligence and *deconfounder*_*N*_ the deconfounder for neuroticsm following the procedure described in Equation 2 and Equation 3.

For supplementary analysis in Figure 2 – Figure supplement 4, proxies and targets were analyzed simultaneously.

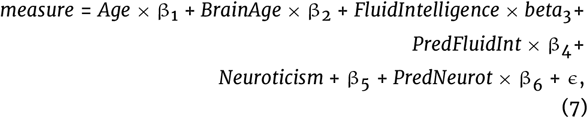

### Software

Preprocessing and model building were carried out using Python 3.7. The NiLearn library was used for processing MRI inputs [100]. We used the *scikit-learn* library for machine learning [109]. For statistical modeling and visualization we used the R-language [110] (version 3.5.3) and its ecosystem: data.table for high-performance manipulation of tabular data, ggplot [111, 112] for visualization and the arm package for parametric bootstrapping [113]. All data analysis code is shared on GitHub [68].

## Availability of source code and requirements

- Project name: “empirical_proxy_measures”
- Project home page: [68]
- Operating system(s): e.g. Platform independent
- Programming language: e.g. Python and R
- Other requirements: e.g. Python 3.6.8 or higher, R 3.4.3 or higher
- License: BSD-3

## Availability of supporting data and materials

Aggregated data supporting the results and figures of this article is available through the GigaScience Database [114] and the “empirical_proxy_measures” code repository [68]. In the future, the individual-level proxy measures obtained from the prediction models in this work will be shared in agreement with the UK Biobank regulations. Please revisit the code repository “empirical_proxy_measures” [68] for details. The input data is available for other researchers via UKBB’s controlled access scheme [115]. The procedure to apply for access [116] requires registering with the UK Biobank and compiling an application form detailing:

- A summary of the planned research
- The UK Biobank data fields required for the project
- A description of derivatives (data, variables) generated by the project

## Declarations

### Author’s Contributions (alphabetic order)

- **Conceptualization**: BT, DB, DE, GV, JH
- **Data curation**: DB, KD
- **Software**: BT, DE, GV, KD
- **Formal analysis**: DE, GV, KD
- **Supervision**: BT, DE, GV
- **Funding acquisition**: GV, JH
- **Validation**: DE, KD
- **Investigation**: DE, KD
- **Visualization**: DE, GV, KD
- **Methodology**: BT, DE, GV
- **Project administration**: DE, GV
- **Writing - original draft**: DE, KD
- **Writing - review and editing**: DB, BT, DE, GV, JH, KD

## Acknowledgements

We would like to thank Dr. Stefania de Vito and Dr. Benjamin de Haas for the critical review and helpful discussion of previous versions of the manuscript. We would like to thank Dr. Julien Dubois and Prof. Ralph Adolphs for helpful discussions in the course of this research project.

## Supporting Information

## Appendix 1: Additional results

**Figure 1 – Figure supplement 1.**
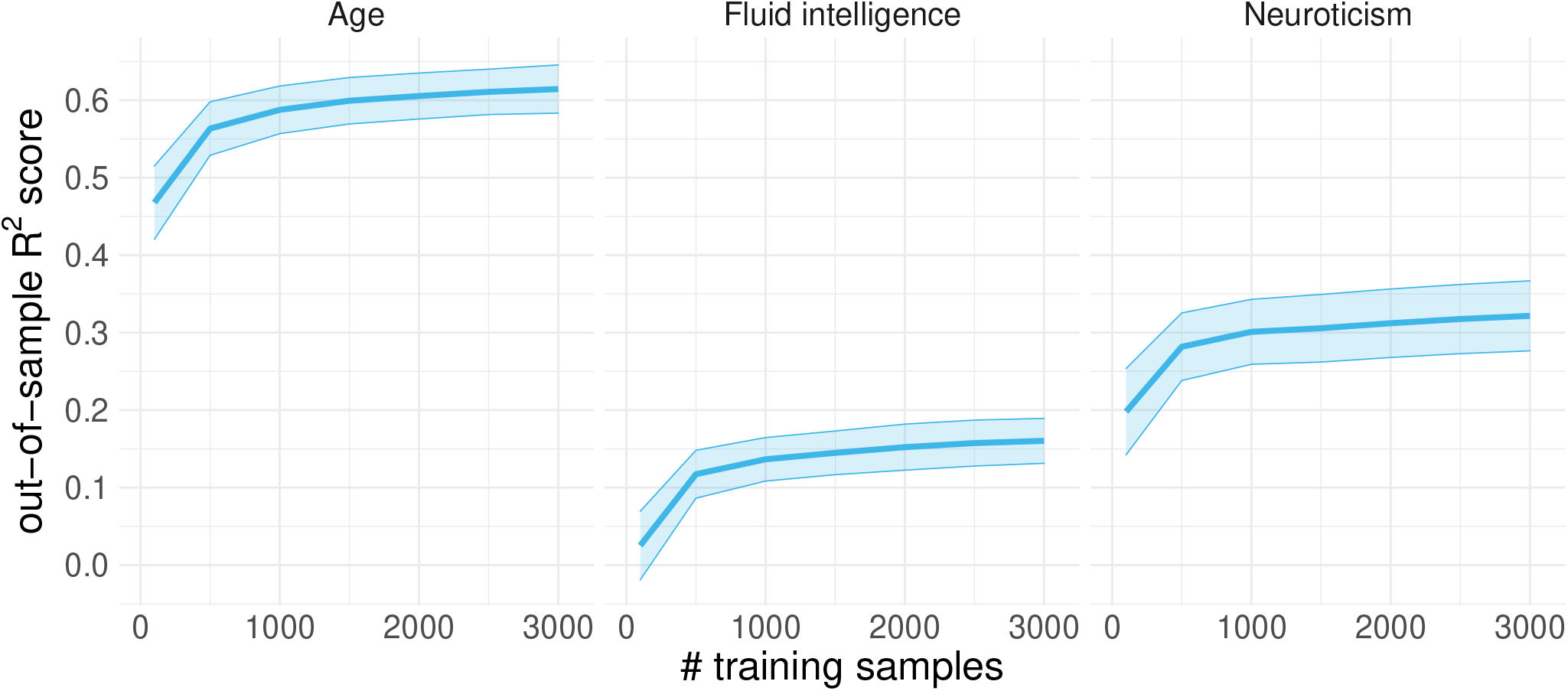
Learning curves on the random split-half validation used for model building. To facilitate comparisons, we evaluated predictions of age, fluid intelligence and neuroticism from a complete set of socio-demographic variables without brain imaging using the coefficient of determination *R*^2^ metric (y-axis) to compare results obtained from 100 to 3000 training samples (x-axis). The cross-validation (CV) distribution was obtained from 100 Monte Carlo splits. Across targets, performance started to plateau after around 1000 training samples with scores virtually identical to the final model used in subsequent analyses. These benchmarks suggest that inclusion of additional training samples would not have led to substantial improvements in performance.

**Figure 2 – Figure supplement 1.**
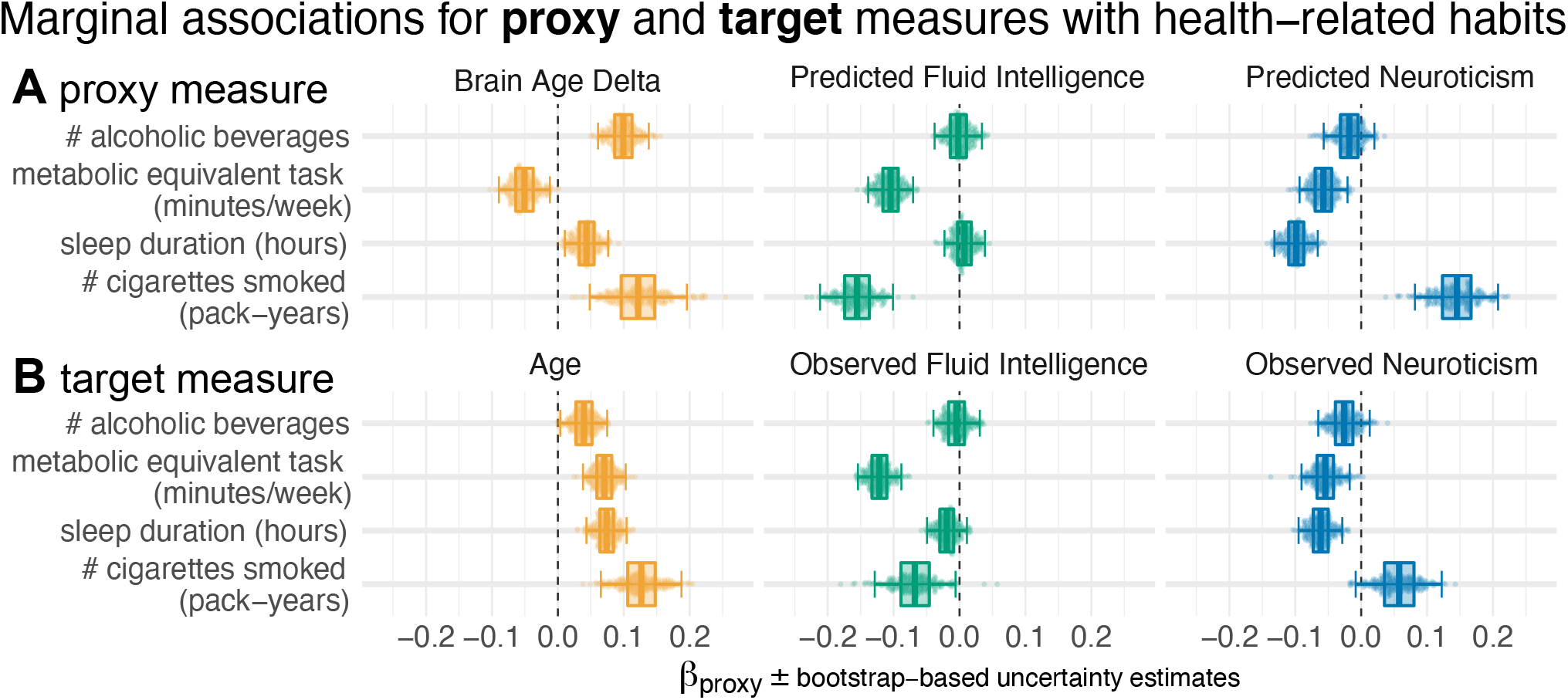
Marginal associations between proxy measures and health-related habits. Marginal (instead of conditional) estimates using univariate regression. Same visual conventions as in Figure 2 – Figure supplement 1.

**Figure 2 – Figure supplement 2.**
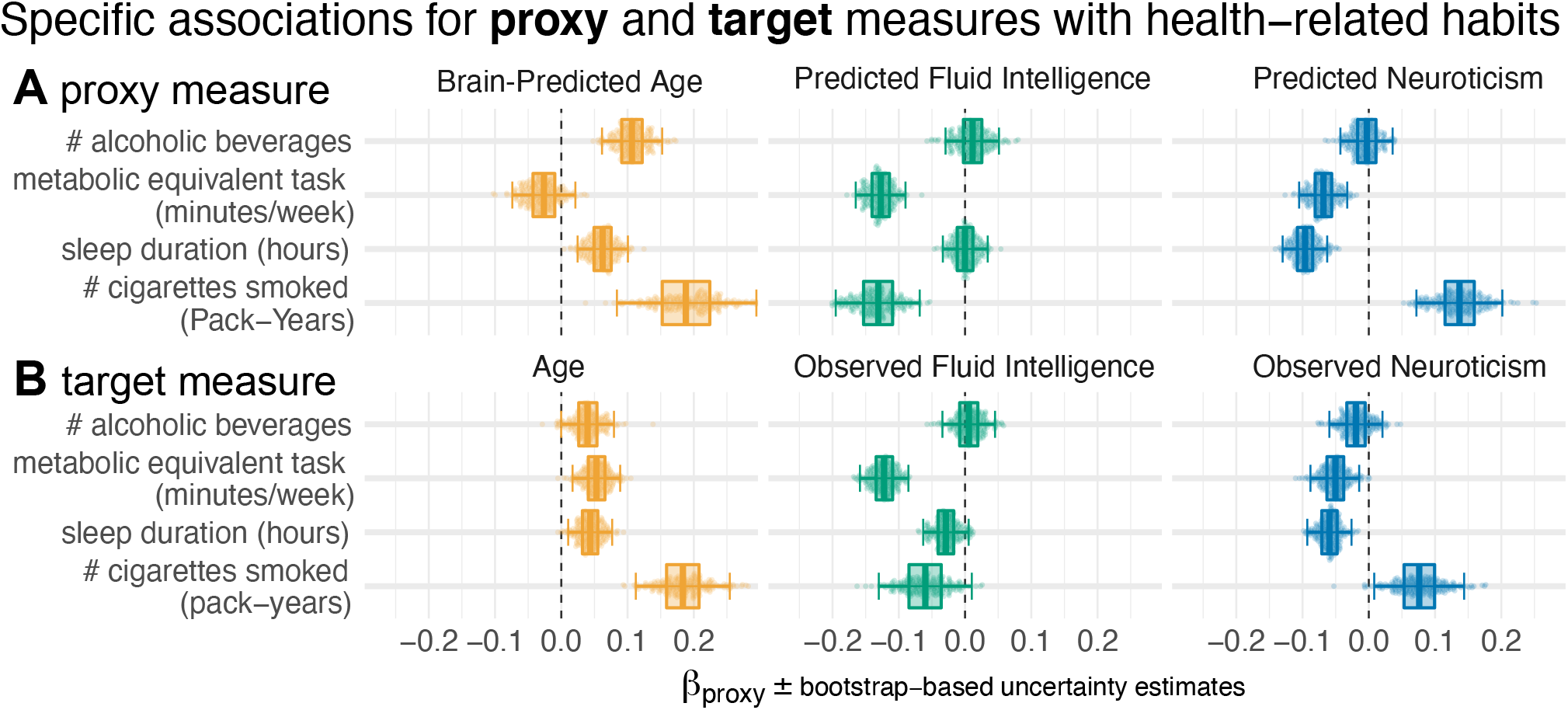
Conditional associations between proxy measures and health-related habits without explicit brain age delta. Conditional estimates using multivariate regression. Instead of the brain age delta, the brain-predicted age is included alongside an age-deconfounder as used in the main analysis. Same visual conventions as in Figure 2.

**Figure 2 – Figure supplement 3.**
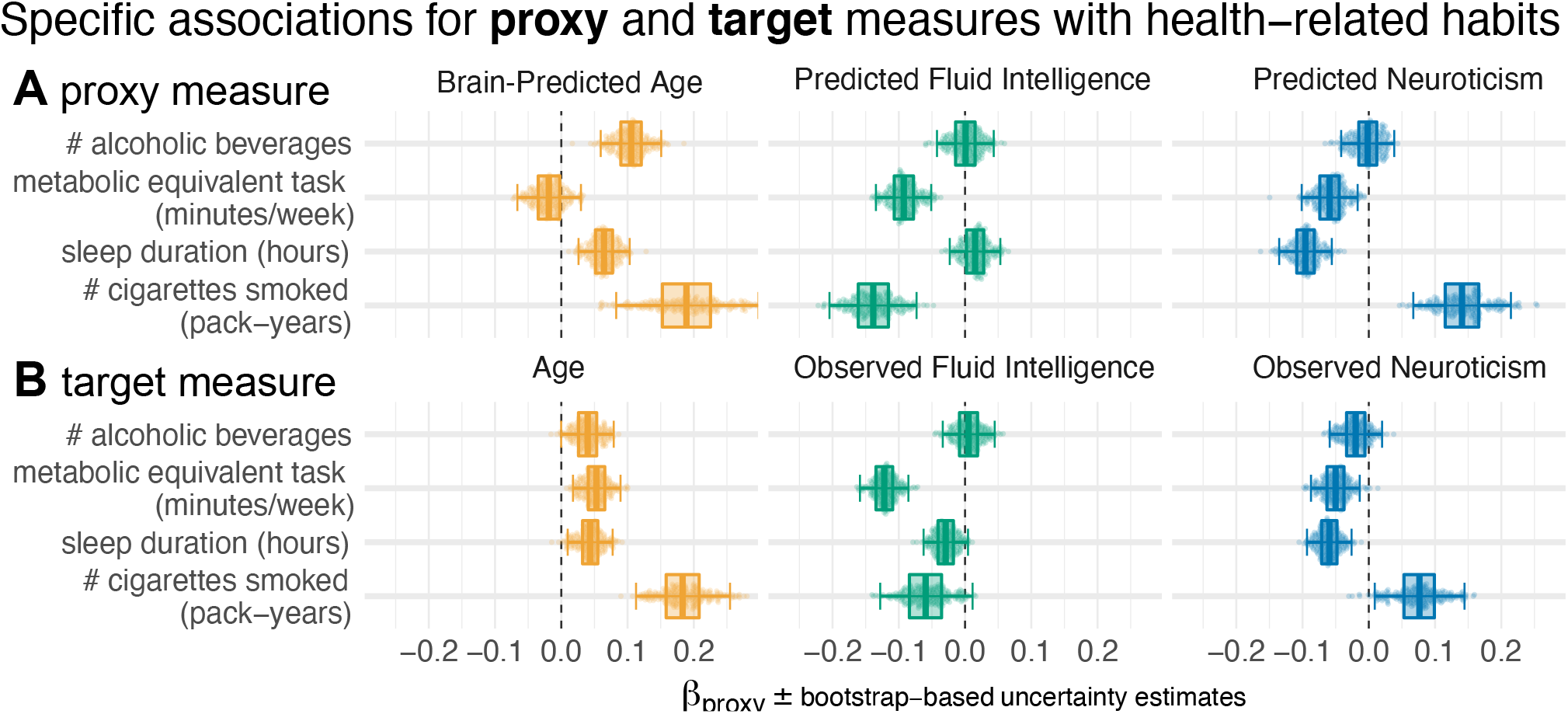
Conditional associations between proxy measures and health-related habits with-proxy-specific deconfounding. Conditional estimates using multivariate regression. Instead of the brain age delta, the brain-predicted age is included alongside an age-deconfounder as used in the main analysis. Moreover, predicted fluid intelligence and neuroticism are deconfounded for the target values at training time, analogous to the brain age predictions. Same visual conventions as in Figure 2.

**Figure 2 – Figure supplement 4.**
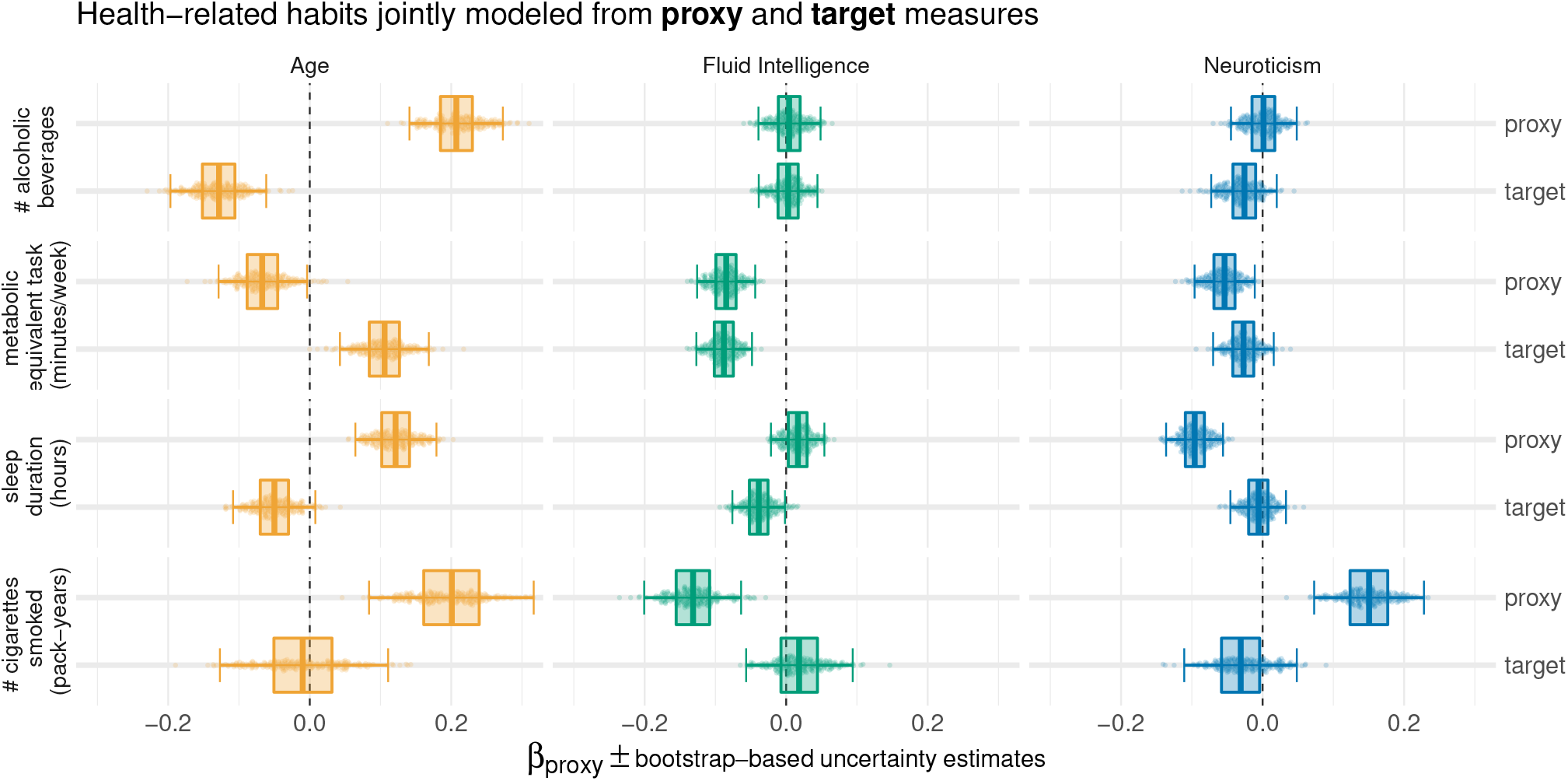
Joint modeling of health-related habits from proxy and target measures. Conditional estimates using multivariate regression. Every health-related habit (double rows) is modeled simultaneously from multiple proxies and targets. Same visual conventions as in Figure 2. Across health-habits, additive effects emerged not only for proxies and targets within the same measure (*e*.*g*. age) but also across measures (*e*.*g*. age and fluid intelligence). For illustration, we shall consider two examples. Regarding alcohol consumption, age was the most important measure and opposite conditional effects were observed for the proxy and the target: Across the age range, people with higher brain age tended to drink more and across the brain-age range, older people tended to drink less. For smoking, the proxy measures were the most important variables with clear non-zero coefficients, pointing in different directions across target domains. Holding fluid intelligence and neuroticism constant (targets and proxies), people with higher brain age tended to have been smoking for a longer time. At the same time, those who scored lower on predicted fluid intelligence across the entire range of age, predicted age, measured fluid intelligence, predicted neuroticism and neuroticism, have been smoking for a longer time. Finally, those who scored higher on predicted neuroticism tended to smoke more across the ranges of all other measures.

**Table S1.**
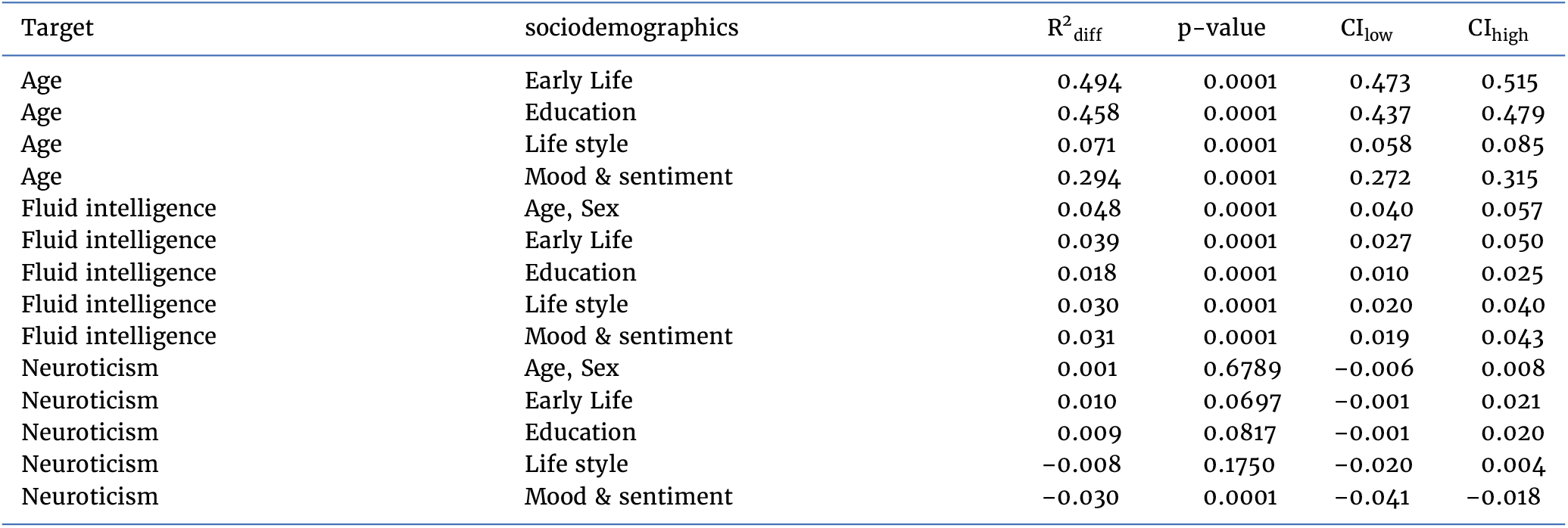
Paired difference between purely sociodemographic and models including brain imaging on held-out data.

| Target | sociodemographics | $R^2_{\text{diff}}$ | p-value | $CI_{\text{low}}$ | $CI_{\text{high}}$ |
| --- | --- | --- | --- | --- | --- |
| Age | Early Life | 0.494 | 0.0001 | 0.473 | 0.515 |
| Age | Education | 0.458 | 0.0001 | 0.437 | 0.479 |
| Age | Life style | 0.071 | 0.0001 | 0.058 | 0.085 |
| Age | Mood & sentiment | 0.294 | 0.0001 | 0.272 | 0.315 |
| Fluid intelligence | Age, Sex | 0.048 | 0.0001 | 0.040 | 0.057 |
| Fluid intelligence | Early Life | 0.039 | 0.0001 | 0.027 | 0.050 |
| Fluid intelligence | Education | 0.018 | 0.0001 | 0.010 | 0.025 |
| Fluid intelligence | Life style | 0.030 | 0.0001 | 0.020 | 0.040 |
| Fluid intelligence | Mood & sentiment | 0.031 | 0.0001 | 0.019 | 0.043 |
| Neuroticism | Age, Sex | 0.001 | 0.6789 | -0.006 | 0.008 |
| Neuroticism | Early Life | 0.010 | 0.0697 | -0.001 | 0.021 |
| Neuroticism | Education | 0.009 | 0.0817 | -0.001 | 0.020 |
| Neuroticism | Life style | -0.008 | 0.1750 | -0.020 | 0.004 |
| Neuroticism | Mood & sentiment | -0.030 | 0.0001 | -0.041 | -0.018 |

**Table S2.** Difference statistics for classification on the held-out set for sociodemographic vs combined approximation.

| Target | $AUC_{\text{diff observed}}$ | p-value | $CI_{\text{low}}$ | $CI_{\text{high}}$ |
| --- | --- | --- | --- | --- |
| Age | 0.013 | 0.0008 | 0.006 | 0.021 |
| Fluid intelligence | -0.031 | 0.0001 | -0.044 | -0.017 |
| Neuroticism | -0.003 | 0.4818 | -0.013 | 0.006 |

**Table S3.** Inferential statistics for joint proxy-target models of health-related habits

|  | Outcome |  |  |  |
| --- | --- | --- | --- | --- |
|  | Alcohol | Activity | Sleep | Smoking |
| predicted Age | 0.208***<br>(0.034) | −0.066**<br>(0.032) | 0.121***<br>(0.029) | 0.200***<br>(0.058) |
| Age | −0.129***<br>(0.035) | 0.105***<br>(0.032) | −0.050*<br>(0.030) | −0.008<br>(0.060) |
| predicted Fluid Intelligence | 0.004<br>(0.022) | −0.085***<br>(0.021) | 0.016<br>(0.019) | −0.132***<br>(0.035) |
| Fluid Intelligence | 0.003<br>(0.022) | −0.088***<br>(0.020) | −0.038**<br>(0.019) | 0.018<br>(0.038) |
| predicted Neuroticism | 0.001<br>(0.024) | −0.054**<br>(0.022) | −0.095***<br>(0.020) | 0.151***<br>(0.040) |
| Neuroticism | −0.026<br>(0.024) | −0.027<br>(0.022) | −0.006<br>(0.020) | −0.031<br>(0.041) |
| Constant | −0.001<br>(0.019) | 0.018<br>(0.018) | 0.017<br>(0.017) | −0.052<br>(0.034) |
| Observations | 2,687 | 3,022 | 3,504 | 896 |
| R<sup>2</sup> | 0.016 | 0.031 | 0.020 | 0.071 |
| Adjusted R<sup>2</sup> | 0.014 | 0.029 | 0.018 | 0.064 |
| Residual Std. Error | 1.004 (df = 2680) | 0.997 (df = 3015) | 0.992 (df = 3497) | 0.992 (df = 889) |
| F Statistic | 7.334*** (df = 6; 2680) | 15.854*** (df = 6; 3015) | 11.733*** (df = 6; 3497) | 11.256*** (df = 6; 889) |

Note:
\*p<0.1; \*\*p<0.05; \*\*\*p<0.01

**Table S4.**
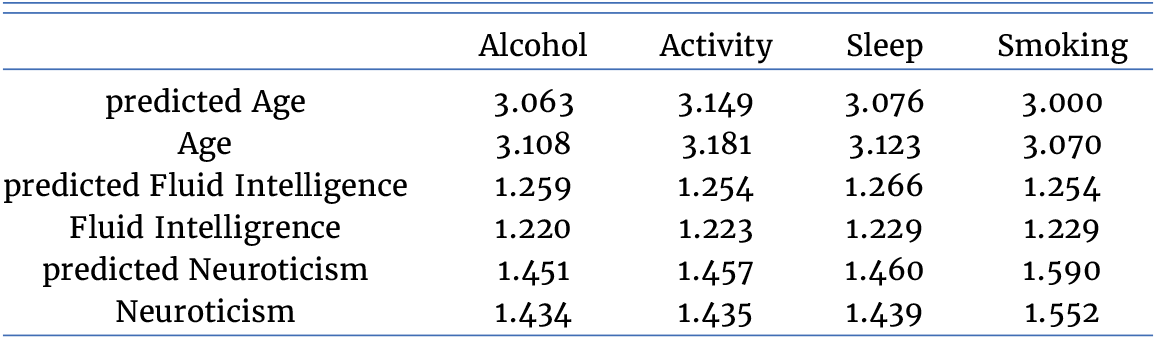
Variance Inflation Factors (VIF) for joint proxy-target models of health-related habits

**Figure 3 – Figure supplement 1.**
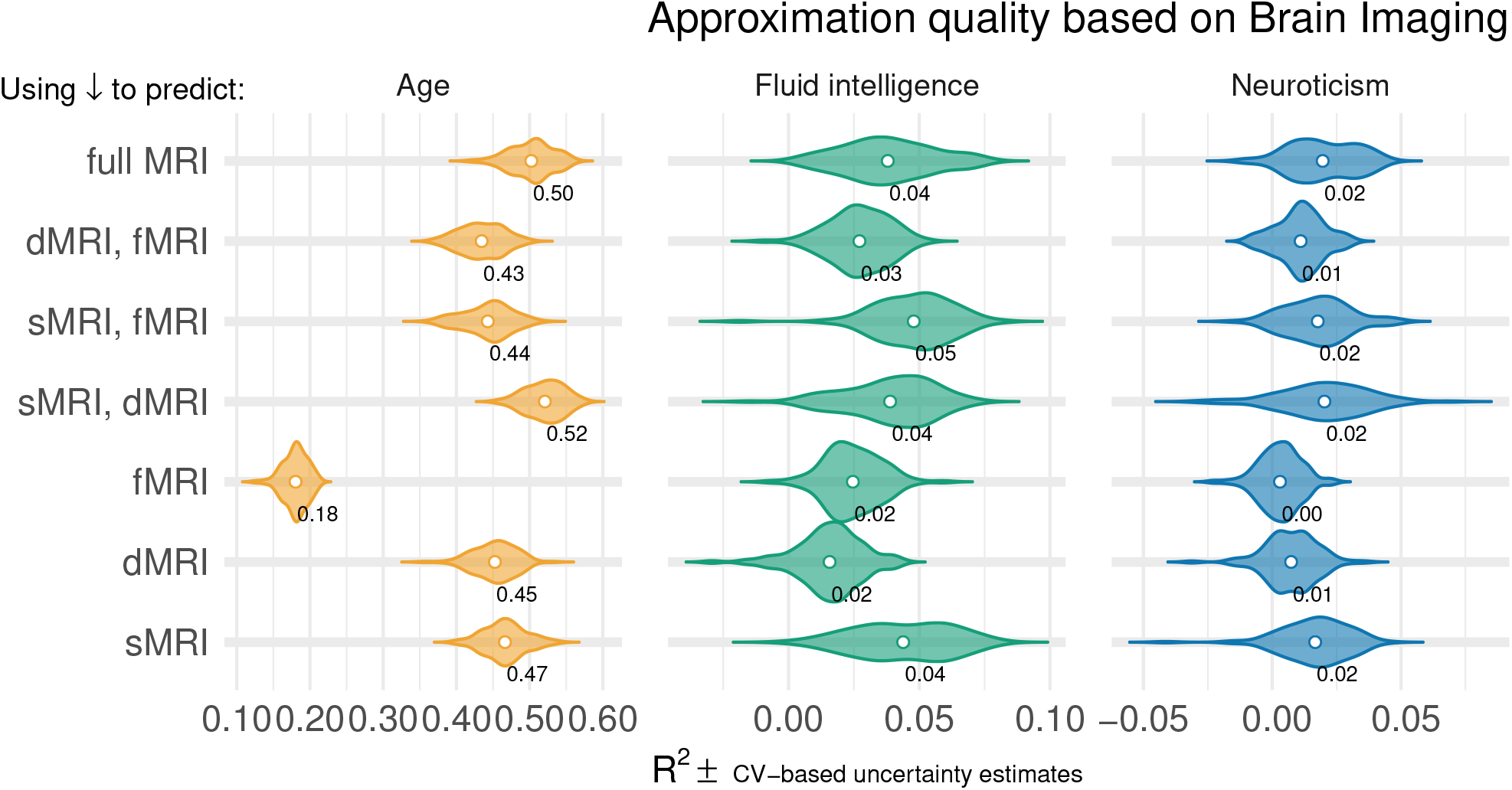
Prediction of individual differences in proxy measures from MRI. Approximation performance using multiple MR modalities on the validation dataset: sMRI, dMRI, rfMRI and their combinations (see Table 1). Visual conventions as in Figure 3. One can see that prediction of age was markedly stronger than prediction of fluid intelligence or prediction of neuroticism. As a general trend, models based on multiple MRI modalities tended to yield better prediction. For simplicity, we based subsequent analyses on the full model based on all MRI data.

**Figure 3 – Figure supplement 2.**
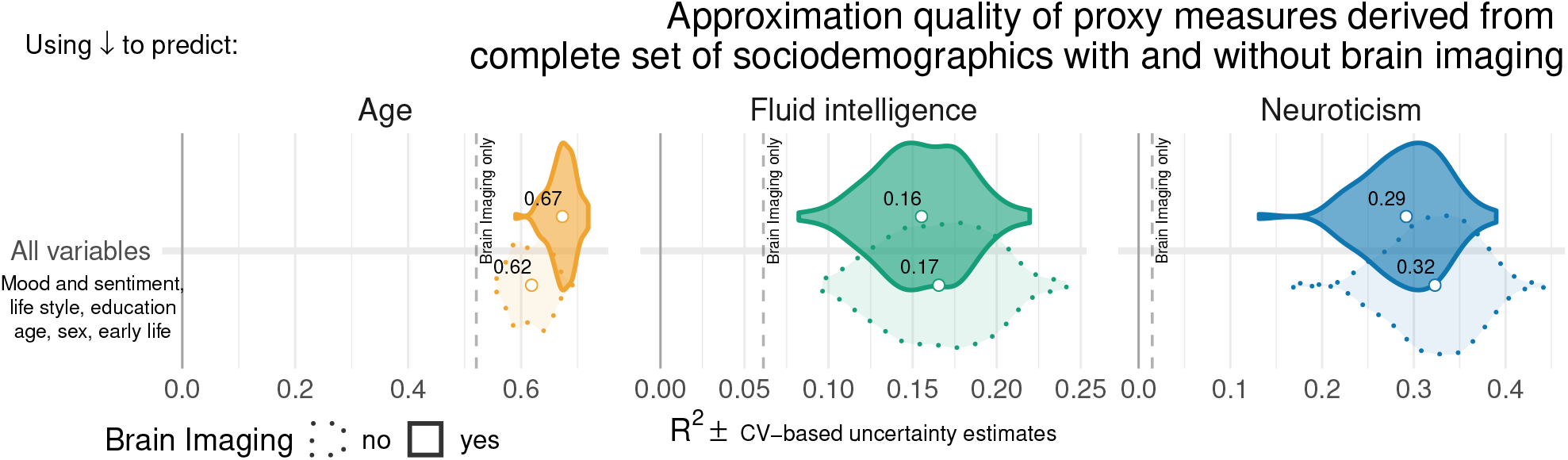
Approximation performance using all sociodemographic data. Approximation performance using all sociodemographic variables with or without brain imaging included on the validation dataset. Visual conventions as in Figure 3. The performance was highly related to the best performing models within each target Figure 3, *i*.*e*., life style for age, education for fluid intelligence and mood & sentiment for neuroticism. This suggests that for each target those specific blocks of predictors were sufficiently explaining the performance. For simplicity, we based subsequent analyses in Figure 4 and Figure 2 on all sociodemographic variables.

**Table S5.** Regression statistics on the held-out set for purely MRI-based approximation.

| Target | $R^2_{\text{observed}}$ | p-value | $CI_{\text{low}}$ | $CI_{\text{high}}$ |
| --- | --- | --- | --- | --- |
| Age | 0.521 | $1 \times 10^{-4}$ | 0.502 | 0.538 |
| Fluid intelligence | 0.061 | $1 \times 10^{-4}$ | 0.052 | 0.070 |
| Neuroticism | 0.015 | $1 \times 10^{-4}$ | 0.005 | 0.024 |

**Table S6.** Classification difference statistics on the held-out set for MRI-based approximation.

| Target | AUC<sub>observed</sub> | p-value | CI<sub>low</sub> | CI<sub>high</sub> |
| --- | --- | --- | --- | --- |
| Neuroticism | 0.590 | $1 \times 10^{-4}$ | 0.566 | 0.614 |
| Age | 0.916 | $1 \times 10^{-4}$ | 0.905 | 0.927 |
| Fluid intelligence | 0.667 | $1 \times 10^{-4}$ | 0.643 | 0.690 |

## Appendix 2: Sociodemographic variables

**Table S7.**
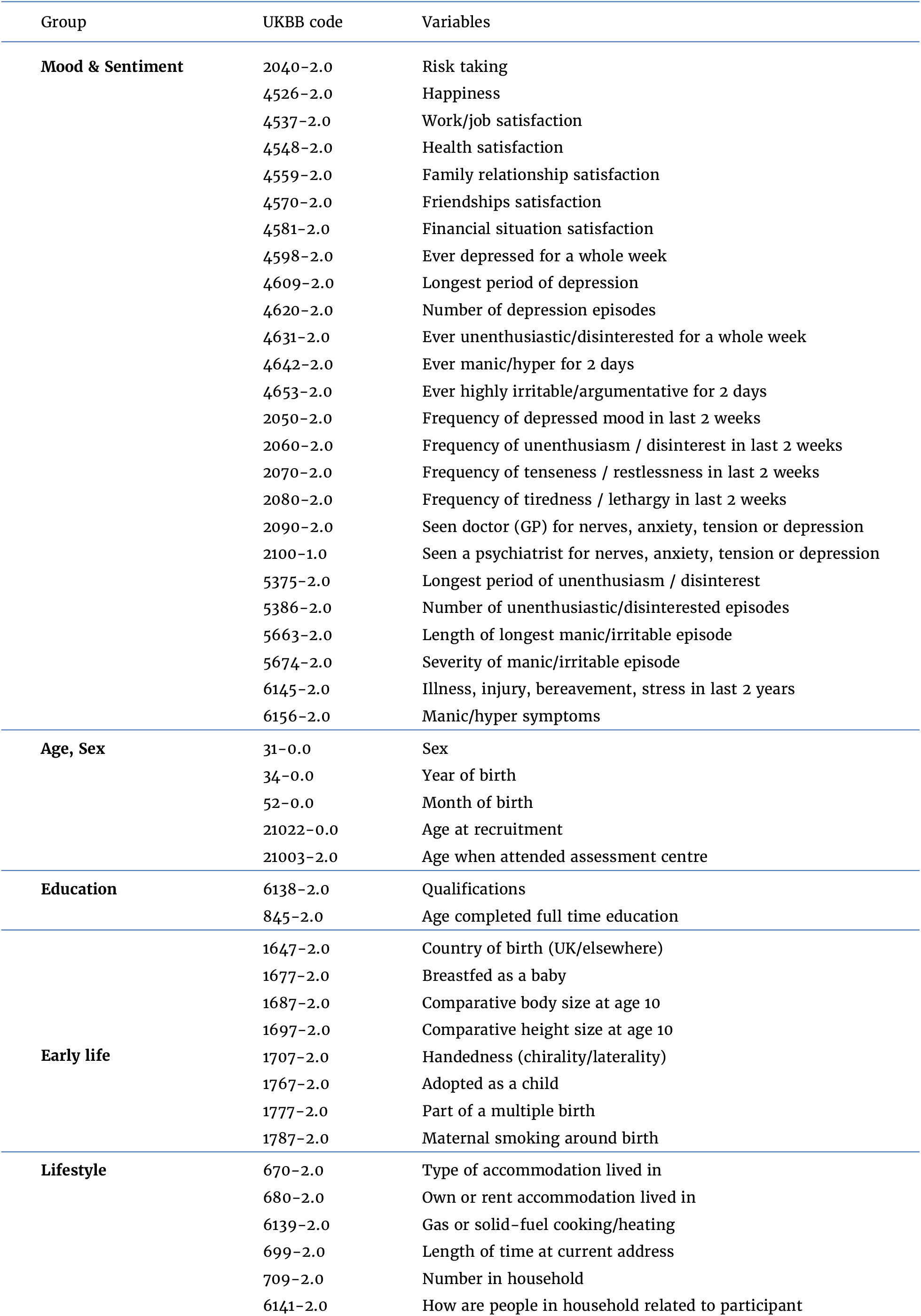
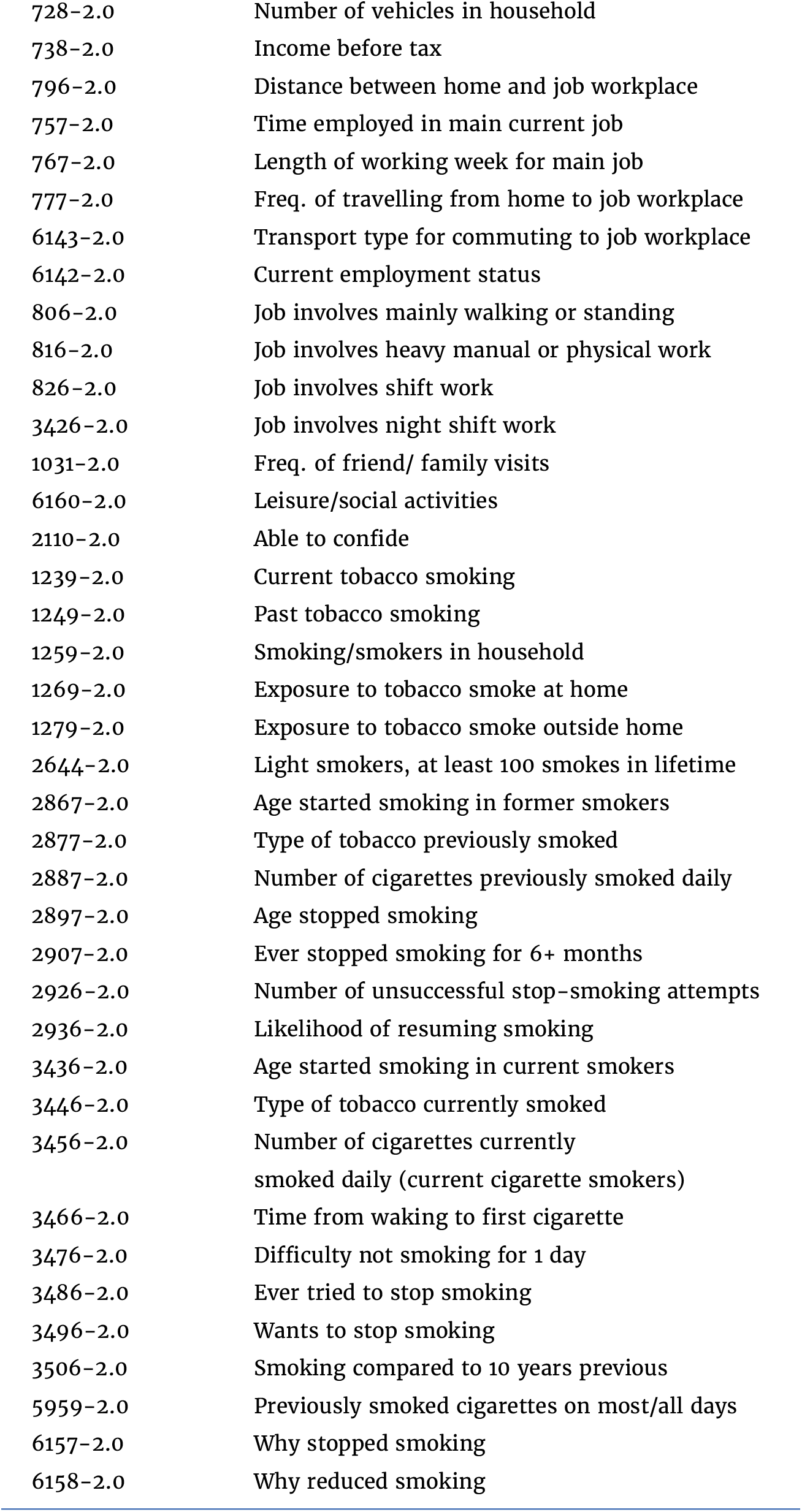
List of variables contained in each block of sociodemographic models: mood & sentiment (MS), Age, Sex (AS), Education (EDU), Early life (EL).

| Group | UKBB code | Variables |
| --- | --- | --- |
| **Mood & Sentiment** | 2040-2.0 | Risk taking |
|  | 4526-2.0 | Happiness |
|  | 4537-2.0 | Work/job satisfaction |
|  | 4548-2.0 | Health satisfaction |
|  | 4559-2.0 | Family relationship satisfaction |
|  | 4570-2.0 | Friendships satisfaction |
|  | 4581-2.0 | Financial situation satisfaction |
|  | 4598-2.0 | Ever depressed for a whole week |
|  | 4609-2.0 | Longest period of depression |
|  | 4620-2.0 | Number of depression episodes |
|  | 4631-2.0 | Ever unenthusiastic/disinterested for a whole week |
|  | 4642-2.0 | Ever manic/hyper for 2 days |
|  | 4653-2.0 | Ever highly irritable/argumentative for 2 days |
|  | 2050-2.0 | Frequency of depressed mood in last 2 weeks |
|  | 2060-2.0 | Frequency of unenthusiasm / disinterest in last 2 weeks |
|  | 2070-2.0 | Frequency of tenseness / restlessness in last 2 weeks |
|  | 2080-2.0 | Frequency of tiredness / lethargy in last 2 weeks |
|  | 2090-2.0 | Seen doctor (GP) for nerves, anxiety, tension or depression |
|  | 2100-1.0 | Seen a psychiatrist for nerves, anxiety, tension or depression |
|  | 5375-2.0 | Longest period of unenthusiasm / disinterest |
|  | 5386-2.0 | Number of unenthusiastic/disinterested episodes |
|  | 5663-2.0 | Length of longest manic/irritable episode |
|  | 5674-2.0 | Severity of manic/irritable episode |
|  | 6145-2.0 | Illness, injury, bereavement, stress in last 2 years |
|  | 6156-2.0 | Manic/hyper symptoms |
| **Age, Sex** | 31-0.0 | Sex |
|  | 34-0.0 | Year of birth |
|  | 52-0.0 | Month of birth |
|  | 21022-0.0 | Age at recruitment |
|  | 21003-2.0 | Age when attended assessment centre |
| **Education** | 6138-2.0 | Qualifications |
|  | 845-2.0 | Age completed full time education |
| **Early life** | 1647-2.0 | Country of birth (UK/elsewhere) |
|  | 1677-2.0 | Breastfed as a baby |
|  | 1687-2.0 | Comparative body size at age 10 |
|  | 1697-2.0 | Comparative height size at age 10 |
|  | 1707-2.0 | Handedness (chirality/laterality) |
|  | 1767-2.0 | Adopted as a child |
|  | 1777-2.0 | Part of a multiple birth |
|  | 1787-2.0 | Maternal smoking around birth |
| **Lifestyle** | 670-2.0 | Type of accommodation lived in |
|  | 680-2.0 | Own or rent accommodation lived in |
|  | 6139-2.0 | Gas or solid-fuel cooking/heating |
|  | 699-2.0 | Length of time at current address |
|  | 709-2.0 | Number in household |
|  | 6141-2.0 | How are people in household related to participant |

*Table S7 continued*
|  |  |
| --- | --- |
| 728-2.0 | Number of vehicles in household |
| 738-2.0 | Income before tax |
| 796-2.0 | Distance between home and job workplace |
| 757-2.0 | Time employed in main current job |
| 767-2.0 | Length of working week for main job |
| 777-2.0 | Freq. of travelling from home to job workplace |
| 6143-2.0 | Transport type for commuting to job workplace |
| 6142-2.0 | Current employment status |
| 806-2.0 | Job involves mainly walking or standing |
| 816-2.0 | Job involves heavy manual or physical work |
| 826-2.0 | Job involves shift work |
| 3426-2.0 | Job involves night shift work |
| 1031-2.0 | Freq. of friend/ family visits |
| 6160-2.0 | Leisure/social activities |
| 2110-2.0 | Able to confide |
| 1239-2.0 | Current tobacco smoking |
| 1249-2.0 | Past tobacco smoking |
| 1259-2.0 | Smoking/smokers in household |
| 1269-2.0 | Exposure to tobacco smoke at home |
| 1279-2.0 | Exposure to tobacco smoke outside home |
| 2644-2.0 | Light smokers, at least 100 smokes in lifetime |
| 2867-2.0 | Age started smoking in former smokers |
| 2877-2.0 | Type of tobacco previously smoked |
| 2887-2.0 | Number of cigarettes previously smoked daily |
| 2897-2.0 | Age stopped smoking |
| 2907-2.0 | Ever stopped smoking for 6+ months |
| 2926-2.0 | Number of unsuccessful stop-smoking attempts |
| 2936-2.0 | Likelihood of resuming smoking |
| 3436-2.0 | Age started smoking in current smokers |
| 3446-2.0 | Type of tobacco currently smoked |
| 3456-2.0 | Number of cigarettes currently<br>smoked daily (current cigarette smokers) |
| 3466-2.0 | Time from waking to first cigarette |
| 3476-2.0 | Difficulty not smoking for 1 day |
| 3486-2.0 | Ever tried to stop smoking |
| 3496-2.0 | Wants to stop smoking |
| 3506-2.0 | Smoking compared to 10 years previous |
| 5959-2.0 | Previously smoked cigarettes on most/all days |
| 6157-2.0 | Why stopped smoking |
| 6158-2.0 | Why reduced smoking |

**Figure S1.**
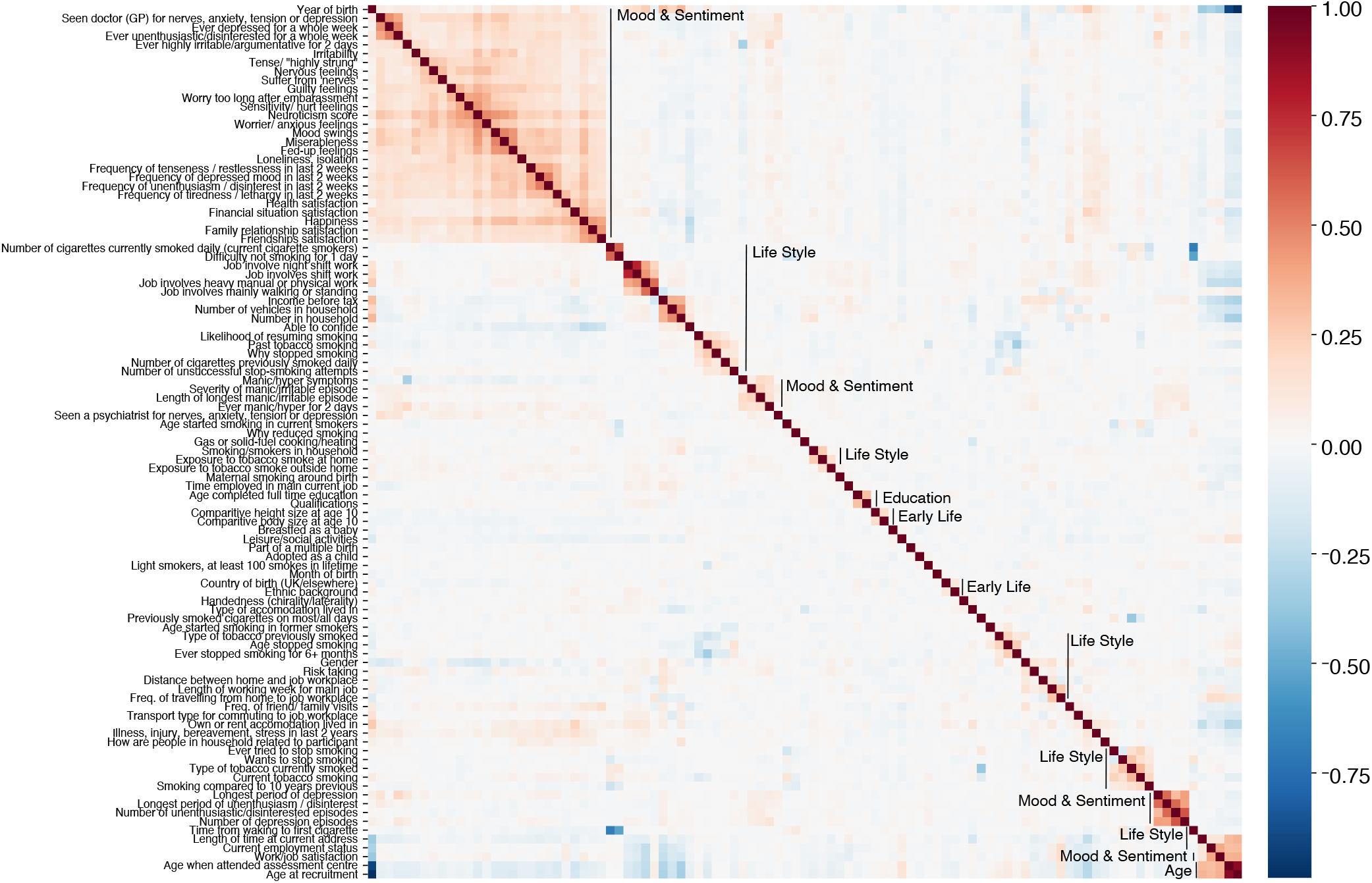
Intercorrelations between sociodemographic inputs. To check the plausibility of the proposed grouping of variables into blocks, we investigated the inter-correlations among the sociodemographic inputs (Table S7). We first applied Yeo-Johnson power transform to the variables yield approximately symmetrical distributions. Then we computed Pearson correlations. One can see that a large majority of variables shows low if any inter-correlations. Strongly inter-correlated blocks emerged, in particular for Mood & Sentiment and Life Style. Note that within the Life Style category many smaller blocks with strong inter-correlation occurred, some of which were obviously related to the circumstance of living such as household or employment status.

## Appendix 3: Impact of Measurement Time

**Figure S2.**
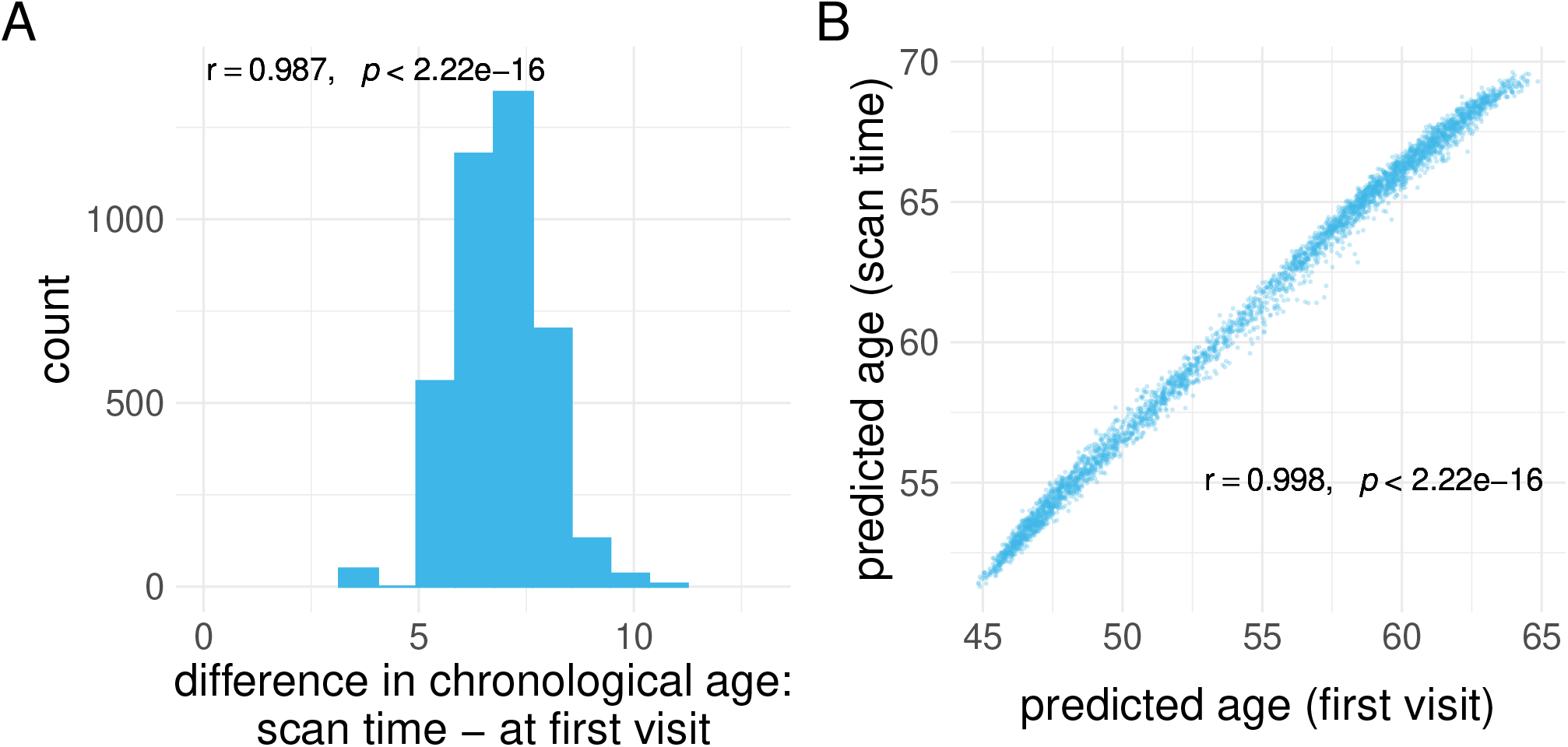
Investigating the age gap between the first visit and the MRI-visit time point. **(A)** Individual gap between age at first visit and MRI-scan time. MRI scans never happened at the first visit, leading to a strictly positive gap greater than five years for most participants. Pearson’s correlation coefficient indicates high rank stability, suggesting that, from a statistical perspective, age at first visit and age at scan time are, essentially, interchangeable. **(B)** Direct comparison of individual-specific age predictions from brain images and sociodemographic data. Same model as in the main analysis (Figure 2). The emerging pattern of association summarized by Pearson’s correlation coefficient suggests that predictions from models either trained on age at the first visit or at MRI-scan time are equivalent.

**Figure S3.**
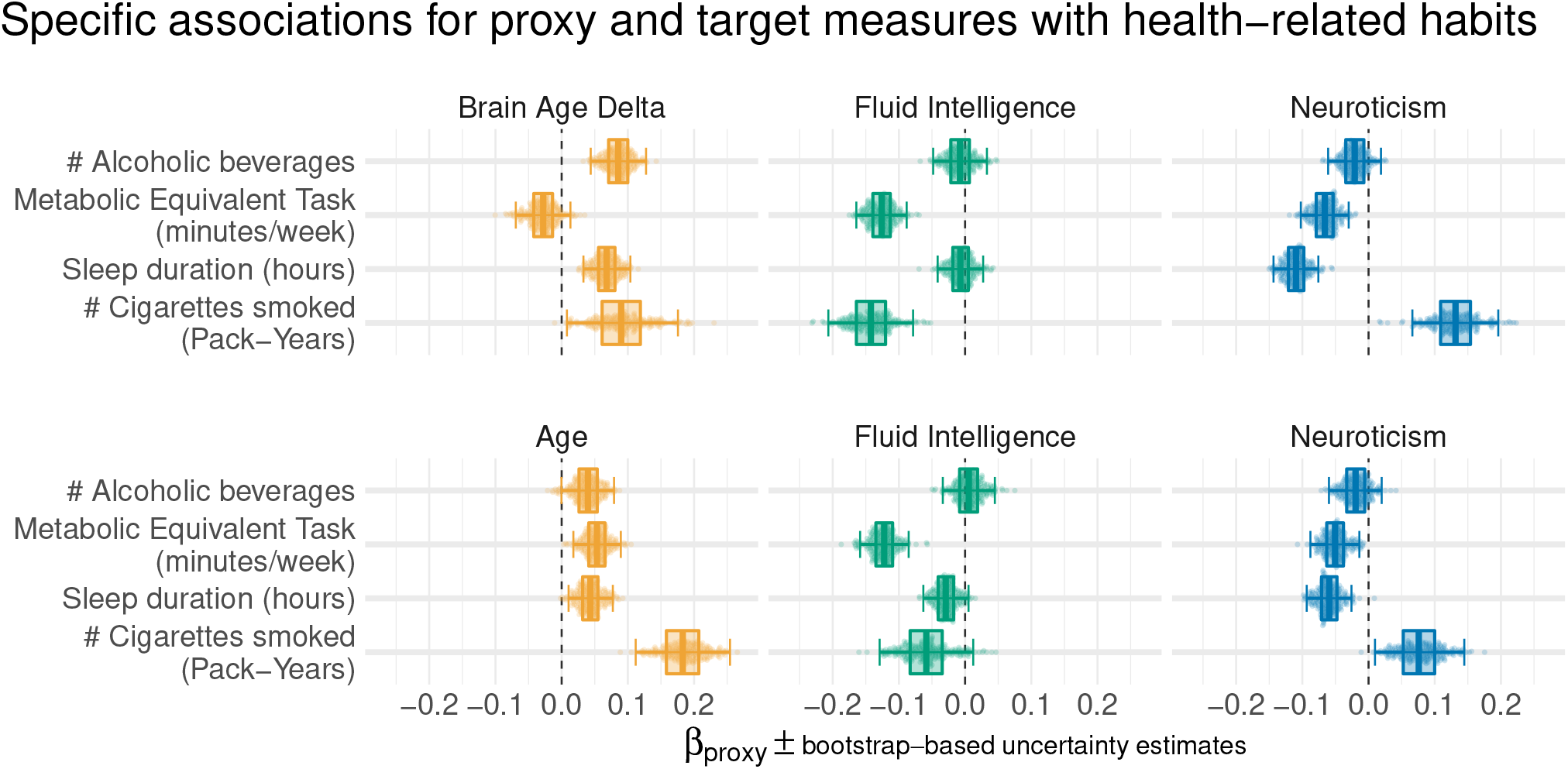
Proxy measures show systematic and complementary out-of-sample associations with health-related habits using age at MRI-scan time. The patterns observed in Figure 2 and global conclusions remain unchanged.

## Footnotes

1 A complete overview of the 13 individual fluid intelligence items is provided by the dedicated user manual [75]

2 For a complete list of Neuroticism questionnaires is provided by the dedicated field descriptions and derivation for variables related to bipolar disorder, major depression status and neuroticism score [76]

3 Regional grey matter volumes [86] Subcortical volumes [87]

4 Diffusion-MRI skeleton measurements [91]

5 We ensured prior to computation that with 100 CV-splits, predictions were available for all subjects.

6 The use of CV-bagging can explain why on figures 3,4, and 3 – Figure supplement 1 the performance was sometimes slightly better on the held-out set compared to the cross-validation on the validation test.

